# Fabrication of Crescent Shaped Microparticles for Particle Templated Droplet Formation

**DOI:** 10.1101/2023.10.06.561257

**Authors:** Yimin Yang, Sergei I. Vagin, Bernhard Rieger, Ghulam Destgeer

## Abstract

Crescent-shaped hydrogel microparticles have been shown to template uniform volume aqueous droplets upon simple mixing with aqueous and oil media for various bioassays. This emerging “lab on a particle” technique requires hydrogel particles with tunable material properties and dimensions. The crescent shape of the particles is attained by aqueous two-phase separation of polymers inside a spherical droplet followed by photopolymerization of the curable precursor. In this work, we have investigated the phase separation of photo-curable poly(ethylene glycol) diacrylate (PEGDA, *M_w_* 700) and dextran (*M_w_* 40,000) for tunable manufacturing of crescent-shaped particles. The particles’ morphology was precisely tuned by following a phase diagram, varying the UV intensity, and adjusting the flow rate ratio between the three streams, containing PEGDA, dextran, and oil, within a microfluidic droplet generation device. The fabricated particles with variable cavity sizes and outer diameters encapsulated uniform aqueous droplets upon mixing with a continuous oil phase. The particles were fluorescently labeled with red and blue emitting dyes at variable concentrations to produce six color-coded particles. The blue fluorescent dye showed a moderate response to the pH change from 1 to 7 in terms of an increase in emitted intensity. The fluorescently labeled particles were able to tolerate an extremely acidic solution (pH 1) but disintegrated within an extremely basic solution (pH 14). The particle-templated droplets were able to effectively retain the disintegrating particle and the fluorescent signal at pH 14, indicating completely segregated compartments.

## 1. Introduction

Biomedical research is rapidly advancing owing to the ability to analyze complex samples at single-cell or single-molecule resolution by high-throughput analytical methods.^1–3^ To achieve high resolution, biological samples are commonly broken into numerous small compartments. Such miniaturized compartments dramatically increase the detectable signals and prevent crosstalk between neighboring compartments.^4–8^ Microwell plates as a commercialized platform have been widely used to perform colorimetric or fluorescent assays in smaller containers. The bottom surface of each well can support cells for growth and provide analyte capture sites for affinity assays.^9–11^ However, expensive benchtop instruments are required to analyze the bioassays within a well.

Recently, droplet microfluidic has been proposed to partition samples.^12–14^ The samples can be compartmentalized into nanoliter to femtoliter droplets rapidly and continuously. Each droplet acts as an isolated reaction compartment for single-cell or molecule detection without crosstalk.^15^ The unique features of droplet microfluidic, such as tiny sample consumption and high throughput, make it a powerful platform for various applications including single-cell analysis^16,17^, nucleic acid detection,^18,19^ enzyme evolution,^20,21^ and drug discovery.^22,23^ However, the standard assay workflow usually needs reagent exchange and washing steps, which can’t directly be achieved in aqueous droplets. In addition, microbeads need to be encapsulated into droplets as affinity capture sites.^24,25^ When the barcoded microbeads are introduced to the droplet, it also allows encoding of each reaction compartment for multiplex detection.^26,27^ However, the encapsulation efficiency of microbeads is dictated by Poisson loading statistics.^28^ The low loading efficiency of microbeads limits the high throughput analysis. Besides, the droplets can’t be prepared ahead of time, and the operation requires specialized microfluidic equipment and skilled personnel.

Recently, particle-templated droplets or *dropicles*, as an emerging compartmentalization method, have demonstrated great potential for numerous biochemical assays in a “lab on a particle” format.^29–35^ The hydrogel particles can be used as a solid phase to support cells or capture biomolecules and as templates to form droplets by agitating with oil simultaneously.^32–35^ The hydrogel particles with a cavity provide a confined region to hold a tiny sample volume or a cell.^30,31,35^ The Poisson loading of cells into particles can be improved by adjusting the cavity size equal to the cell diameter.^29^ Particle-templated droplet formation is a user-friendly strategy that relies only on common lab apparatus such as a pipette or benchtop vortex mixer;^34^ therefore, it eliminates the need for microfluidic platforms for compartmentalization. However, the fabrication of the particles still benefits from microfluidic flow sculpting or uniform droplet generation.

Structured microparticles have been fabricated with different functions and shapes using droplet microfluidics.^36–38^ The crescent-shaped particles have been produced by photopolymerizing a curable precursor within a droplet of an aqueous two-phase system (ATPS).^29,39–43^ One of the most frequently used ATPS contains poly(ethylene glycol) diacrylate (PEGDA) and dextran as two components with a range of different molecular weights (*M_w_*). The ATPS droplets separate into two immiscible phases as the concentrations of the individual components increase beyond a critical value, of which the photocurable precursor is cross-linked by UV light. In addition to the polymer precursor concentrations in the aqueous medium, the separation of ATPS is also affected by the polymer *M_w_*, and sample temperature.^44,45^ The physicochemical properties of cured particles, such as their size, structure, stiffness, porosity, etc., critically depend on the starting ATPS components. Therefore, it’s essential to investigate the controllable fabrication of crescent-shaped particles with different ATPS combinations. An ATPS combination (PEGDA, *M_w_* 700, and Dextran, *M_w_* 40,000), which we have investigated in this work, has been used previously to fabricate crescent-shaped particles; however, a thorough characterization of this combination using a phase diagram was missing.^43^ Moreover, the existing crescent-shaped particles have demonstrated little barcoding ability, which is a desired feature for multiplex assays.

In this work, we have studied the phase separation of PEGDA (*M_w_* 700) and dextran (*M_w_* 40,000) to fabricate crescent-shaped particles in a controllable manner by using a droplet microfluidic platform (**Figure 1**a-d). Core-shell droplets were formed by flowing aqueous PEGDA and aqueous dextran solutions through a Y-junction of the microfluidic device as the dispersed phase and a fluorocarbon oil with a surfactant through the flow-focusing segment of the microfluidic device as the continuous phase. As the PEGDA phase separated from dextran within the segmented droplet, the PEGDA shell was crosslinked in the presence of a biocompatible photoinitiator (PI) upon exposure to UV light to cure solid crescent-shaped particles. We have systematically investigated the effect of UV intensity, flow rates of aqueous and oil streams, and concentrations of polymer precursors on the curability and dimensions of crescent-shaped particles. We separately introduced blue emitting (MA-P4VB)^46–48^ and red-emitting (MA-RhB) dyes (see **Figure S8** for the dye structures)^49^ into the cured PEGDA networks. This was achieved by photo-copolymerization at different dye concentrations and resulted in covalent bonding of the dye molecules to color-code multiple crescent-shaped particles. The fluorescent crescent-shaped particles with different dimensions were used as templates to generate isolated dropicles upon simple mixing with aqueous and oil phases using a common lab pipette for multiplexed detection (Figure 1e). In addition, we have also studied the effects of solution pH, varied from 1 to 14, on the fluorescent particles within a bulk solution and inside the dropicles.

**Figure 1.**
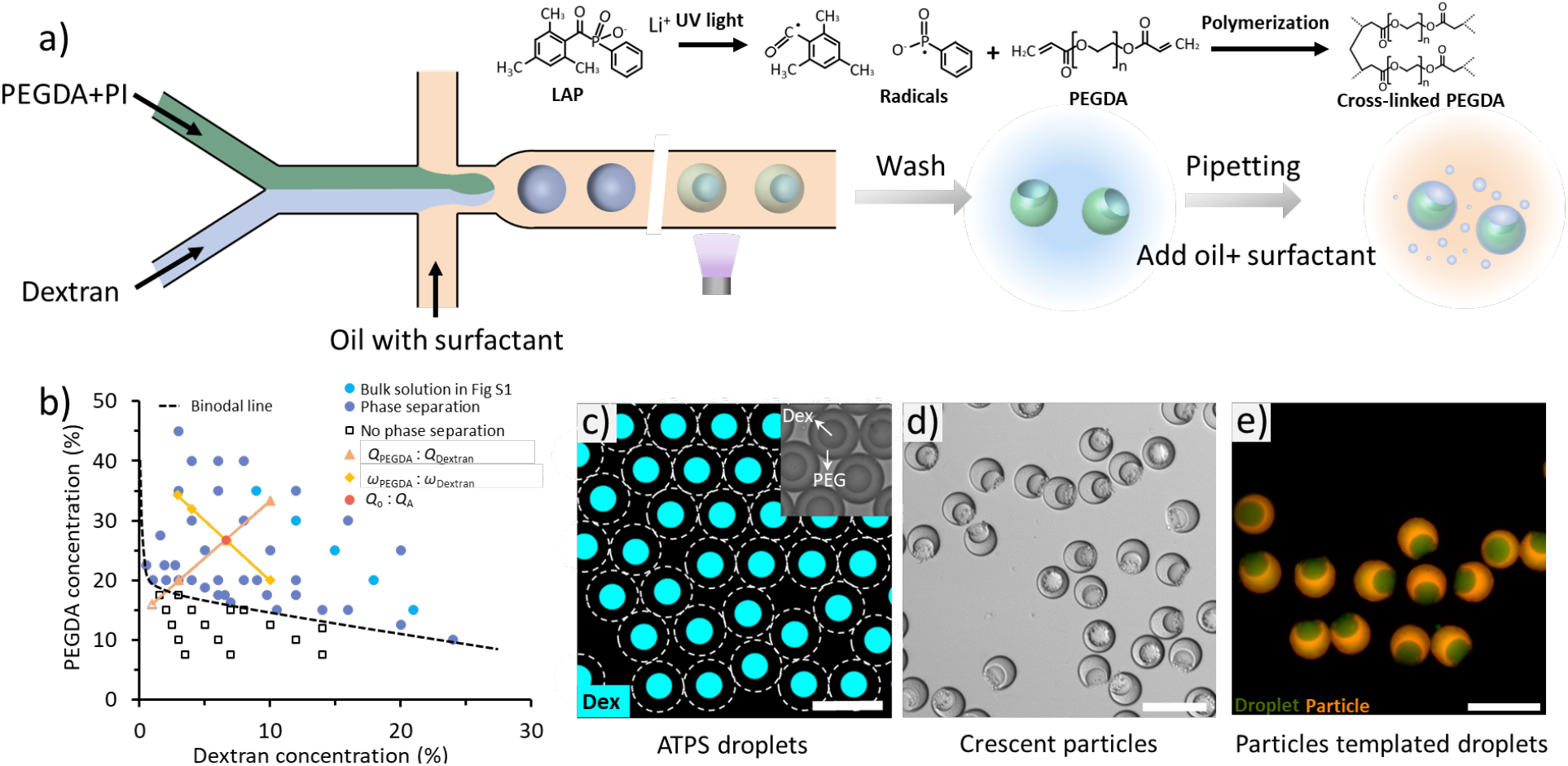
Crescent-shaped particle fabrication and particle-templated droplet (*dropicle*) formation. (a) Crescent-shaped particles were produced using a Y-junction microfluidic device. An aqueous solution comprised of PEGDA and photoinitiator was co-flowed with a solution containing dextran in the microfluidic device. The co-flow was emulsified at the cross-junction by the outer continuous oil phase containing fluorinated Novec 7500 with 0.5% surfactant. (b) Experimentally determined phase diagram of PEGDA (*M_w_* 700) and dextran (*M_w_* 40,000). (c) After droplet formation, the PEGDA and dextran phase-separated, resulting in dextran-rich core and PEGDA-rich shell droplets. The dextran phase was mixed with fluorescein-labeled dextran for visualization purpose. The droplets were exposed to UV light at the end of the device to crosslink the PEGDA shell. (d) The cured particles within droplets were collected, then emulsions were broken, and oil and dextran were removed by a washing step. (e) Particle-templated droplets were formed by pipetting. Fluorescence microscope image of dropicles formed with red-emitting fluorescently labeled particles and green-emitting fluorescein solution. Scale bars are 100 μm.

## 2. Result and Discussion

### 2.1. Phase separation of PEGDA/dextran system

A typical ATPS phase diagram is divided into miscible and immiscible areas by a binodal line defining the critical concentrations of the two dissolved polymers that would clearly phase separate above the binodal curve (Figure 1b).^44,45,50^ To draw the phase diagram of PEGDA (*M_w_* 700) and dextran (*M_w_* 40,000) before emulsification using a droplet generator, we analyzed the phase separation behavior in a bulk within a tube. A series of samples with varying concentrations of PEGDA and dextran were prepared and mixed at varied volume ratios (**Table S1**). Then the mixtures were vortexed and centrifuged to induce phase separation. For example, the five samples in **Figure S1** were prepared by mixing aqueous solutions of 50% w/w PEGDA and 30% w/w dextran in volume ratios of 7:3, 3:2, 1:1, 2:3, and 3:7. A line separating the PEGDA rich (top) and dextran rich (bottom) phases is visible in all five samples which indicate that phase separation occurred in these bulk solutions. A relatively higher density of dextran allowed it to sink to the bottom of the low-density PEGDA phase at equilibrium.^45^ The corresponding effective concentrations are labeled as phase separation points in Figure 1b.

### 2.2. Continuous microfluidic fabrication of crescent-shaped particles

Based on phase separation experiments in bulk, we designed and fabricated a Y-junction microfluidic device to form ATPS droplets. PEGDA with photoinitiator LAP and dextran were injected into the device through separate inlets as dispersed fluids. The co-flow was emulsified into water/oil (w/o) droplets by an outer continuous oil fluid comprised of Novec 7500 and 0.5% w/w fluorinated surfactant. The typical flow rates of PEGDA, dextran, and continuous flow streams were 50, 25, and 1500 μL/h, respectively. Subsequently, PEGDA and dextran in droplets rapidly phase separated on-chip to form dextran-core/PEGDA-shell droplets after ∼3 s as the concentration of PEGDA and dextran in the droplet was above the binodal line (Figure 1c). The morphology of the ATPS droplet depends on the balance of interfacial energies between the PEGDA-rich and dextran-rich phases with the outer oil phase. The dextran-rich phase was more hydrophilic than the PEGDA-rich phase, thus, dextran prefers to interact with water than the outer oil phase to minimize interfacial energies. Similarly, a relatively hydrophobic PEGDA-rich phase within the droplet prefers to form a shell close to the oil-water interface that surrounds the dextran-rich phase at the core.^45,51^ The PEGDA shell was crosslinked as the droplets flowed past an on-chip UV light exposure window close to the outlet. The crescent-shaped particles were collected by breaking the emulsion and washing them with hexane, ethanol, and water (Figure 1d). By contrast, when the concentration of PEGDA and dextran in the droplet was below the binodal line, spherical particles were obtained due to the complete miscibility of PEGDA and dextran. For the fabrication of crescent-shaped particles, the concentrations of PEGDA and dextran were chosen above the binodal line.

### 2.3. Crescent-shaped particles morphology modulation

#### 2.3.1 The effect of UV intensity

The UV intensity directly affects the polymerization of particles. To investigate how the UV intensity influences the crosslink of PEGDA, we varied the UV intensity from 25% to 100% of the nominal power of the LED light source connected to the microscope to fabricate crescent-shaped particles (**Figure 2**a, b). The concentration of the photoinitiator (PI) remained constant at 1% w/w. The flow rates of the PEGDA phase, dextran phase, and oil phase were kept constant at 50 μL/h, 25 μL/h, and 1500 μL/h, respectively. When the UV intensity increased from 25% to 75%, the outer diameter of particles (*D*_p_) gradually increased from 43.0 μm to 54.5 μm on average. A further increase in intensity to 100% resulted in a minor change in *D*_p_ to 56.3 μm (Figure 2d). It indicated that the particles were almost fully crosslinked at a UV intensity of 75%. The cavity diameter of the cured particles (*D*_c_) showed little change as the UV intensity was increased. The on-chip polymerization enabled the phase-separated droplets to be crosslinked immediately and prevented downstream coalescence. The particles crosslinked at lower intensities still showed structural integrity and good uniformity with the coefficient of variance (CV) values less than 4.0% after the washing step. In addition, we observed in the scanning electron microscopy (SEM) images that the brim of the particle cavities developed a rough texture as the particles crosslinked at high UV intensity (>75%) (Figure 2b). We hypothesized that the roughness was caused by a partially crosslinked thin PEGDA layer between dextran and oil phases. Most of that thin PEGDA layer would have washed away along with the dextran inner phase; however, an under-crosslinked PEGDA segment at the brim of the cavity remained to cause a rough texture. Here, we used a UV intensity of 100% to cure particles for further experiments. We also evaluated the swelling behavior of cured particles upon rehydration from a dried state (**Figure S2**a,b). The dried hydrogel particles absorbed water rapidly and reached swelling equilibrium states in approximately 2 minutes. As the particles got swollen by taking up water, the *D*_p_ increased from 36.3 μm to 54.4 μm during rehydration.

**Figure 2.**
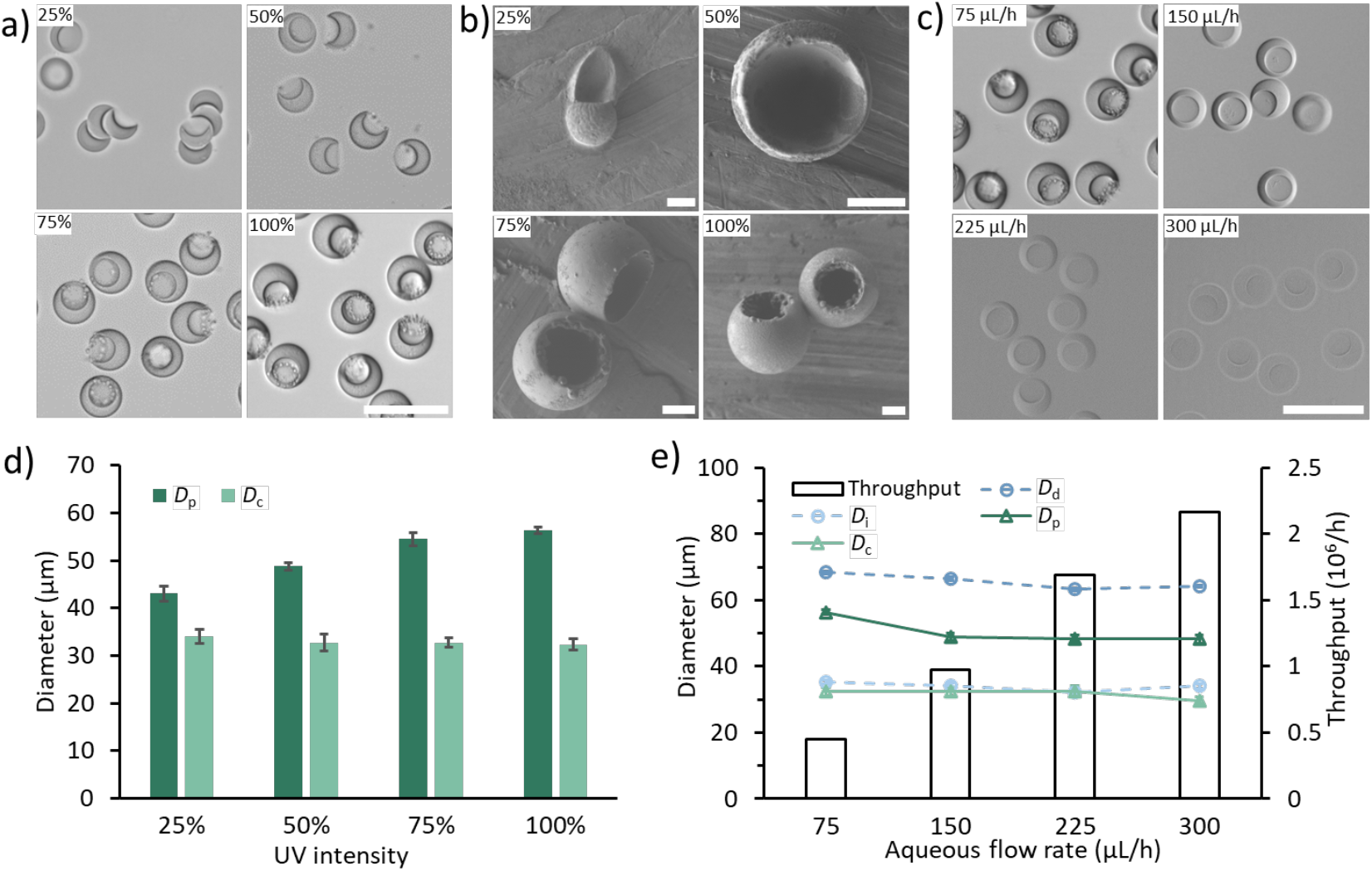
Crescent-shaped particles fabricated with variable UV intensity and flow rates. Microscope (a) and SEM (b) images of crescent-shaped particles fabricated with four different exposure intensities (25%-100%), and their outer diameter (*D*_p_), and cavity diameter (*D*_c_) after washing (d). (c) Microscope images of crescent-shaped particles fabricated with four different aqueous flow rates (75-300 μL/h). (e) The droplet and particle dimensions as a function of aqueous flow rate. The variation in the outer and cavity diameter of the particle (*D*_p_, *D*_c_) and the outer and inner diameter of the droplet (*D*_d_, *D*_i_) with the fabrication throughput. The scale bars are 100 μm in images a, c, and 10 μm in image b.

To observe the throughput of particle fabrication, we increased the total flow rate of three fluids with a maximum UV intensity exposure (Figure 2c,e). The concentration of the PI was kept constant at 1%w/w. We fixed the flow rate ratio of the continuous oil phase to the dispersed aqueous phase at 20, and the flow rate ratio of the PEGDA stream to the dextran stream at 2 to keep droplet morphology the same. The droplet dimensions only changed slightly during the increasing flow rate (**Figure S3**). For example, the inner and outer diameters of droplets (*D*_i_ and *D*_d_) were measured as 35.3 μm and 68.4 μm, respectively, when the total flow rate of the aqueous stream (PEGDA + dextran) was 75 μL/h. After the aqueous flow rate increased to 300 μL/h, *D*_i_ and *D*_d_ only slightly decreased to 34.0 μm and 64.2 μm, respectively. Typically, the UV exposure time of the precursors was reduced as the flow rate increased. As a result, the *D*_p_ decreased from 56.3 μm to 48.2 μm when the aqueous flow rate increased from 75 μL/h to 300 μL/h (Figure 2e). The visibility of particles also gradually decreased due to the lower crosslinking density (Figure 2c). However, because the dextran was uncurable, the *D*_c_ remained constant around 31.8 μm which was close to *D*_i_. Moreover, at each aqueous flow rate, the *D*_p_ was smaller than the *D*_d_ because of oxygen inhibition at the water-oil interface.^52,53^ As calculated from the mass balance, a throughput of 2.16×10^6^ particles per hour can theoretically be achieved while maintaining a reasonable monodispersity of cured particles (Figure 2e).

#### 2.3.2 Flow rate ratio of PEGDA and dextran

The crescent-shaped particle dimensions can be controlled by adjusting the concentration of PEGDA and dextran within the aqueous droplet by varying the flow rate ratio of PEGDA and dextran (*Q*_PEGDA_:*Q*_Dextran_) on-chip (**Figure 3**a-c). In a typical experiment, we used 40% w/w PEGDA and 20% w/w dextran as dispersed fluid and varied the value of *Q*_PEGDA_:*Q*_Dextran_. Other fabrication parameters, such as the total flow rate of dispersed fluid and continuous oil fluid, and the UV intensity were kept constant. The microscope images of droplets and particles at different flow rate ratios are shown in Figure 3a. As *Q*_PEGDA_:*Q*_Dextran_ increased from 1:1 to 6:1, the actual in-droplet concentration of PEGDA increased from 20% to 34.3%, and that of dextran decreased from 10% to 2.8%. These concentration combinations were still lying within the phase separation area of the phase diagram (Figure 1b). Thus, the PEGDA and dextran underwent phase separation in droplets and formed core-shell droplets. The resulting ATPS droplets were highly uniform in their size. For instance, when *Q*_PEGDA_:*Q*_Dextran_ was 2:1, the average *D*_d_ and *D*_i_ were 68.4 μm (CV=0.6%) and 35.3 μm (CV=1.6%), respectively (Figure 3c). The volume ratios of PEGDA-shell and dextran-core (*V*_PEGDA_:*V*_Dextran_) linearly increased from 2.5 to 20.3 as the *Q*_PEGDA_:*Q*_Dextran_ increased (**Figure S4**). After phase-separated droplets were crosslinked by UV light, crescent-shaped particles with different cavity sizes were obtained. Thereby, the cavity size was flexibly tuned with respect to the particle by adjusting *Q*_PEGDA_:*Q*_Dextran_. With increasing *Q*_PEGDA_:*Q*_Dextran_ from 1:1 to 6:1, the relative cavity diameter of particle *R*_c_ (*D*_c_/*D*_p_) decreases from 74.0% to 38.6% (Figure 3b). The cured particles also showed narrow size distributions and good monodispersities. The *D*_p_ and *D*_c_ are 56.6 μm with a 0.8% CV and 35.0 μm with a 3.1% CV, respectively, as *Q*_PEGDA_:*Q*_Dextran_ was 2 (Figure 3c). Due to the flow rates of aqueous and oil fluids being fixed, the *D*_d_ was nearly unchanged. However, the *D*_p_ showed a minor change owing to the different concentrations of PEGDA and photoinitiator in ATPS droplets at equilibrium while exposed to UV light. Specifically, the particles showed the largest variation in the relative change of outer diameter Δ*D* ((*D*_d_-*D*_p_)/*D*_p_) when the *Q*_PEGDA_:*Q*_Dextran_ was 1 because of the lowest actual concentration of photoinitiator and PEGDA in the droplet (Figure 3b).

**Figure 3.**
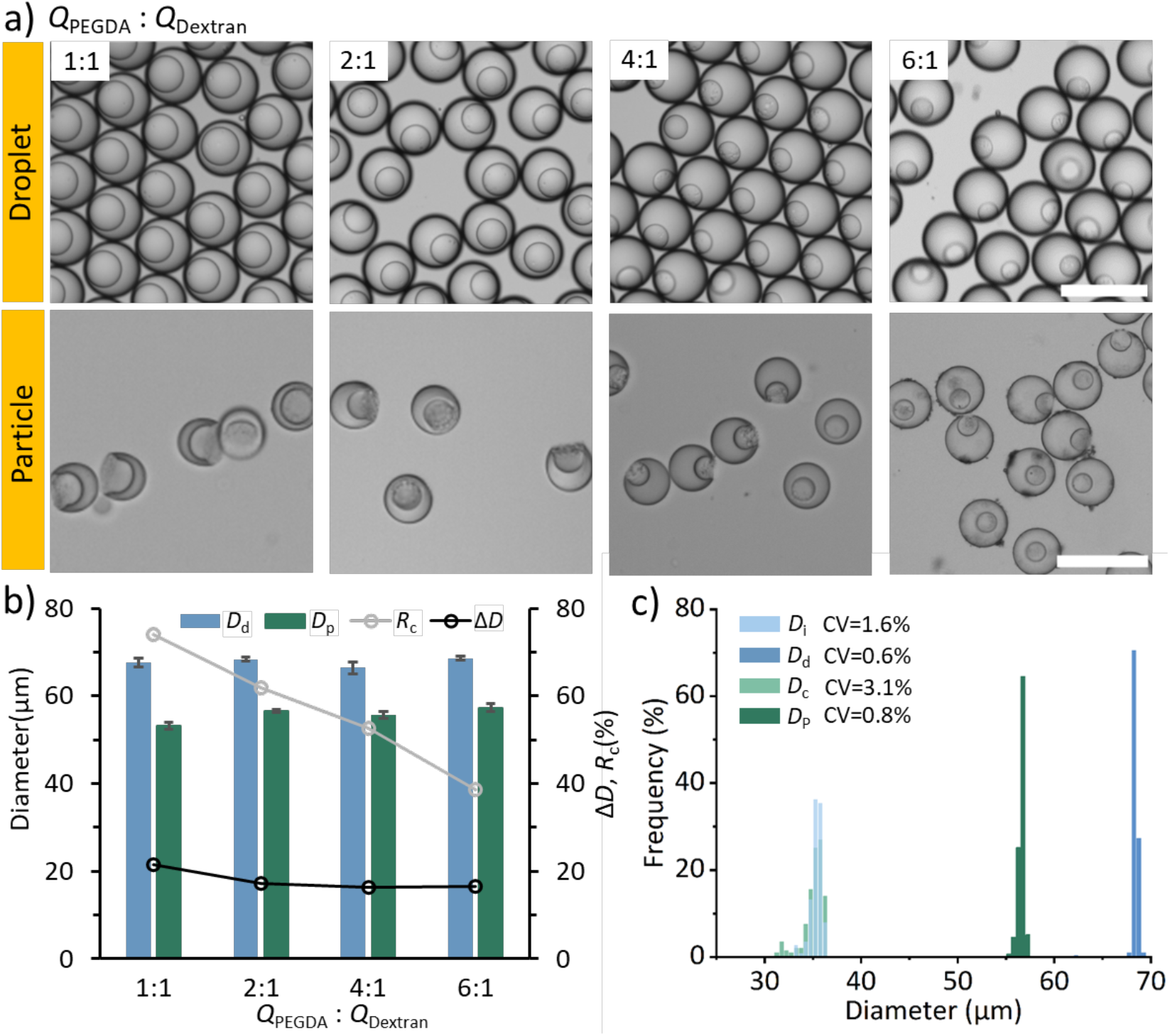
Particle shape was tuned by varying the flow rate ratio of PEGDA and dextran (*Q*_PEGDA_: *Q*_Dextran_) on the chip. (a) Microscope images of droplets and particles with four different *Q*_PEGDA_: *Q*_Dextran_ from 1:1 to 6:1. (b) Effects of *Q*_PEGDA_: *Q*_Dextran_ on the value of *D*_p_, *D*_d_, the relative cavity diameter of particle *R*_c_ (*D*_c_ / *D*_p_), and the relative change of outer diameter Δ*D* ((*D*_d_-*D*_p_)/*D*_p_)). (c) The Histogram of core-shell droplets and crescent-shaped particles as *Q*_PEGDA_: *Q*_Dextran_ was 2. The scale bars represent 100 μm.

#### 2.3.3 Concentration of PEGDA and dextran in bulk

The modulation of particle morphology can also be achieved by directly tuning the concentrations of PEGDA and dextran in bulk (stock) solutions to adjust the concentrations of PEGDA and dextran within the droplet (**Figure 4**a,b). Typically, we randomly choose four groups of PEGDA and dextran with different bulk concentrations to fabricate particles (A: PEGDA 32% w/w, dextran 2% w/w; B: PEGDA 40% w/w, 6% dextran w/w; C: PEGDA 40% w/w, dextran 20% w/w; D: PEGDA 50% w/w, dextran 30% w/w). Other experiment parameters were kept constant. The flow rate ratios of PEGDA and dextran were 1 in groups A and B, and 2 in groups C and D. After emulsification, the actual in-droplet concentrations of PEGDA and dextran (*ω*_PEGDA_:*ω*_Dextran_) were 16%:1%, 20%:3%, 26.7%:6.7%, 33.3%:10%. The generated droplets and particles are shown in Figure 4a. When the *ω*_PEGDA_:*ω*_Dextran_ was 16%:1%, the cured particles were spherical in the absence of phase separation (Figure 1e). The in-droplet concentrations of PEGDA and dextran were lower than the binodal line in the phase diagram. Above the binodal line, as the concentrations of the components were increased, the phase separation enabled the fabrication of crescent-shaped particles with varied cavity diameters. The *R*_c_ value varied from 0 to 60.8%. The *D*_d_ was almost the same with different *ω*_PEGDA_:*ω*_Dextran_ due to the fixed flow rates of the aqueous and oil streams. However, the *D*_p_ gradually increased from 42.1 μm to 61.3 μm due to the increasing PEGDA concentration within the droplet at equilibrium. Reversibly, the Δ*D* decreased from 38.4% to 9.7% as more PEGDA was cured within the same droplet volume (Figure 4b).

**Figure 4.**
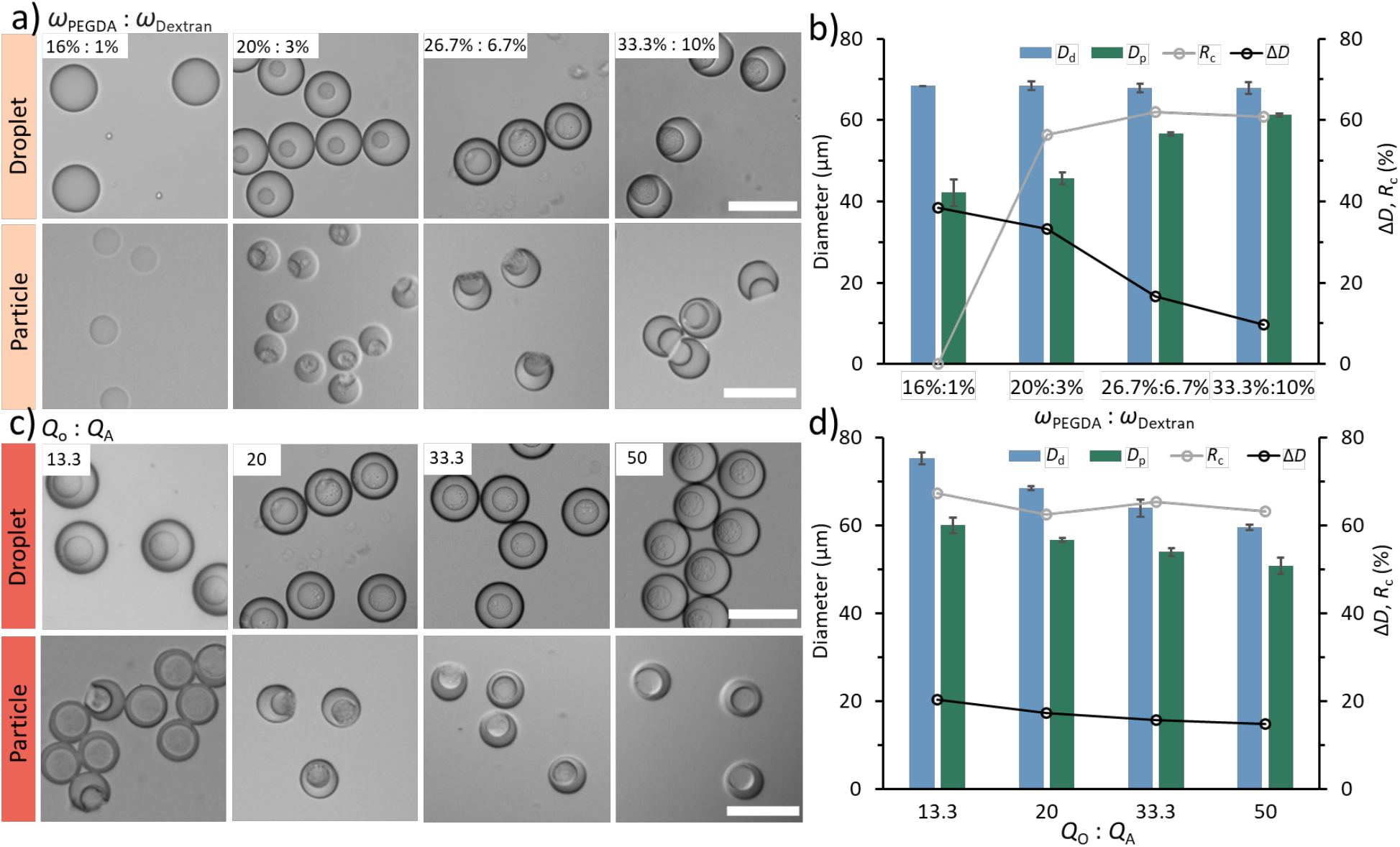
Effect of bulk polymer concentrations and flow rate ratios on particle dimensions. (a, b) Particle shapes were tuned by adjusting the initial polymer concentrations within the bulk sample. (a) Microscope images of droplets and particles with four different mass fractions of PEGDA and dextran (*ω*_PEGDA_: *ω*_Dextran_). (b) Effects of *ω*_PEGDA_: *ω*_Dextran_ on the *D*_p_, *D*_d_, *R*_c_, and Δ*D*. (c, d) Particle shapes were tuned by varying flow rate ratios of continuous to aqueous flow streams (*Q*_O_: *Q*_A_). (c) Microscope images of droplets and particles with four different *Q*_O_: *Q*_A_ (13.3-50). (b) Effects of *Q*_O_: *Q*_A_ on the *D*_p_, *D*_d_, *R*_c_, and Δ*D*. The scale bars represent 100 μm.

#### 2.3.4 Flow rate ratio of continuous and dispersed flow streams

The flow rate ratio of continuous oil and dispersed aqueous flow streams also affected the morphology of cured particles (Figure 4c,d). To obtain particles at different flow rate ratios of continuous and dispersed streams (*Q*_o_:*Q*_A_), we fixed the flow rate ratio of PEGDA to dextran (*Q*_PEGDA_:*Q*_Dextran_) at 2 and varied *Q*_o_:*Q*_A_ from 13.3 to 50 by decreasing the combined flow rate of the aqueous stream. UV intensity was kept constant at 100%. As shown in Figure 4c, with increasing *Q*_o_:*Q*_A_ from 13.3 to 50, the *D*_d_ slowly decreased from 75.3 μm to 59.5 μm. The *D*_p_ and *D*_c_ showed the same decreasing trend. Specifically, *D*_p_ decreases from 60.0 μm to 50.7 μm. Moreover, the *R*_c_ and Δ*D* remained constant at around 64.5% and 17.0%, respectively, since the *Q*_PEGDA_:*Q*_Dextran_ and concentrations of PEGDA and dextran were fixed (Figure 4d). These results indicated that the particle morphology could be easily and precisely controlled by simply adjusting the flow rate of fluids.

### 2.4. Fluorescent encoding of the crescent-shaped particles

To encode the crescent-shaped particles, we added various concentrations of blue-emitting (ex. 395 nm/em. 495 nm) and red-emitting (ex. 548 nm/em. 570 nm) fluorescent dyes possessing methacrylate functional groups to the PEGDA phase (**Figure 5**a,b). The fluorescent monomers were homogeneously distributed within the droplets and had no impact on the phase separation of PEGDA and dextran (**Figure S5**). After the curing and washing steps, the fabricated particles retained the fluorescent colors because the two fluorescent dyes were immobilized in the PEGDA network via chemical covalent bonds due to copolymerization. Respectively, the mean intensity of particles increased with the increasing concentration of fluorescent dyes (Figure 5c). As a proof of concept, we cured six different color-coded particles with three non-overlapping intensities for the two dye colors (blue and red). The six color-coded particles could be distinguished easily, suggesting that they can be used for a multiplexing purpose.

**Figure 5.**
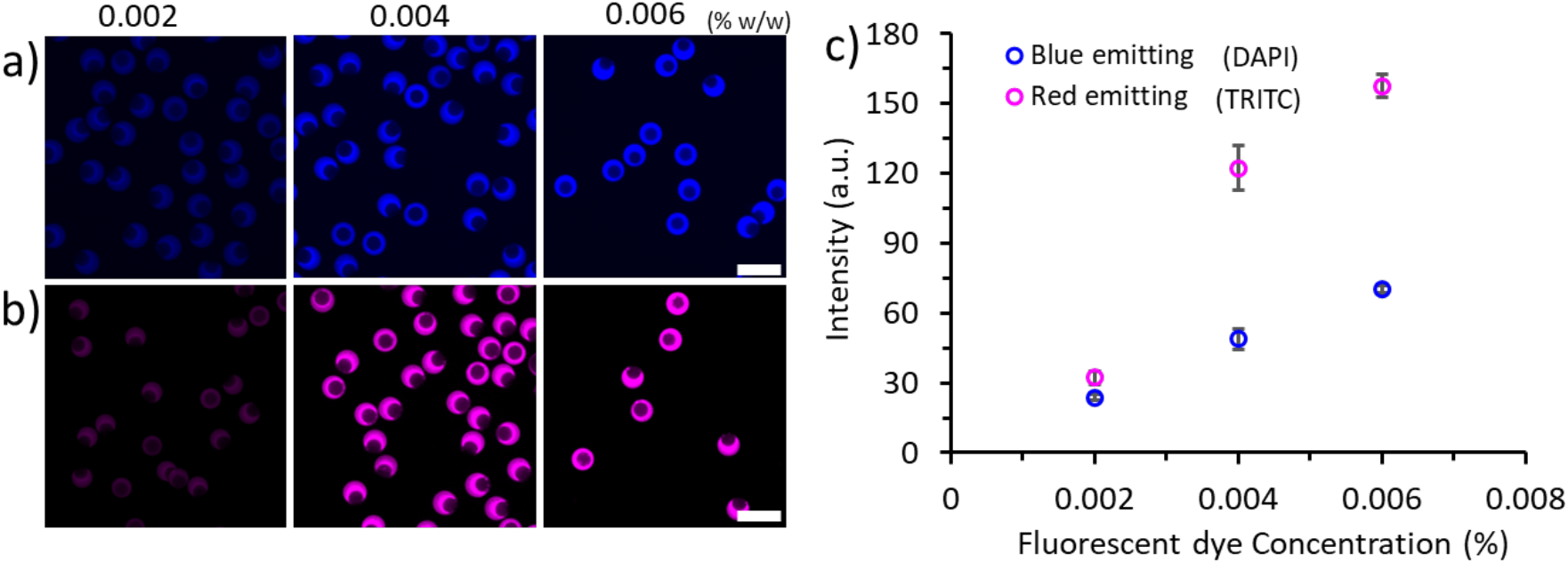
Controllable fabrication of fluorescent crescent-shaped particles. (a, b) Fluorescence microscope images of the particles fabricated with three different concentrations (0.002-0.006w/w %) of two types of fluorescent dyes. (c) The mean fluorescent intensity increased with the concentration of the blue and red fluorescent dyes. The scale bars represent 100 μm.

### 2.5. Particle-templated droplet or dropicles generation

The crescent-shaped particles with tunable size were used as templates to form dropicles (**Figure 6**a-c). The particles suspended in an aqueous solution were mixed with an oil and surfactant solution by simply pipetting. The aqueous volume was split up into numerous tiny compartments holding uniform volume around the solid particles, whereas an even larger number of satellite droplets with much smaller volume was generated in the background that can be easily filtered out during the data analysis. The continuous oil phase around the dropicles ensures isolated compartments of pL-scale volumes with minimal crosstalk between the neighboring compartments. The crescent-shaped particles with variable *R*_c_ were successfully used to form uniform dropicles (CV < 5%). A wide range of encapsulated dropicle volumes was realized, from ∼1.2 pL to ∼0.4 pL, by using particles fabricated with the flowrate ratio of *Q*_PEGDA_:*Q*_Dextran_ from 1:1 to 6:1 (Figure 3 and Figure 6a). The CV of dropicle and corresponding volumes showed a very minor increase with the increasing *R*_c_. For the particles with the largest *R*_c_ of 74.0%, the CV values for diameter and volume were 4.4% and 13.2%, which decreased to 2.2% and 6.6%, respectively, for the smallest *R*_c_ of 38.6% (**Figure S6**). After fluorescent encoding, the crescent particles can serve as barcoded solid phase to compartmentalize biological samples containing multiple targets. To demonstrate the possibility of multiplexed detection, we implemented fluorescently encoded particles with different intensities and fluorescent colors to form dropicles (Figure 6 b,c). The two types of dyes, red and blue colors, were directly identified from the fluorescent color to decode the dropicles (Figure 6b). Particles fabricated with three concentrations (0.002-0.006% w/w) of each dye were mixed to form dropicles (Figure 6c). The fluorescent intensities were measured to identify the dye concentration after distinguishing the type of dye (Figure 6d). Each dropicle could be decoded to obtain the barcode information by combining the fluorescent color and intensity.

**Figure 6.**
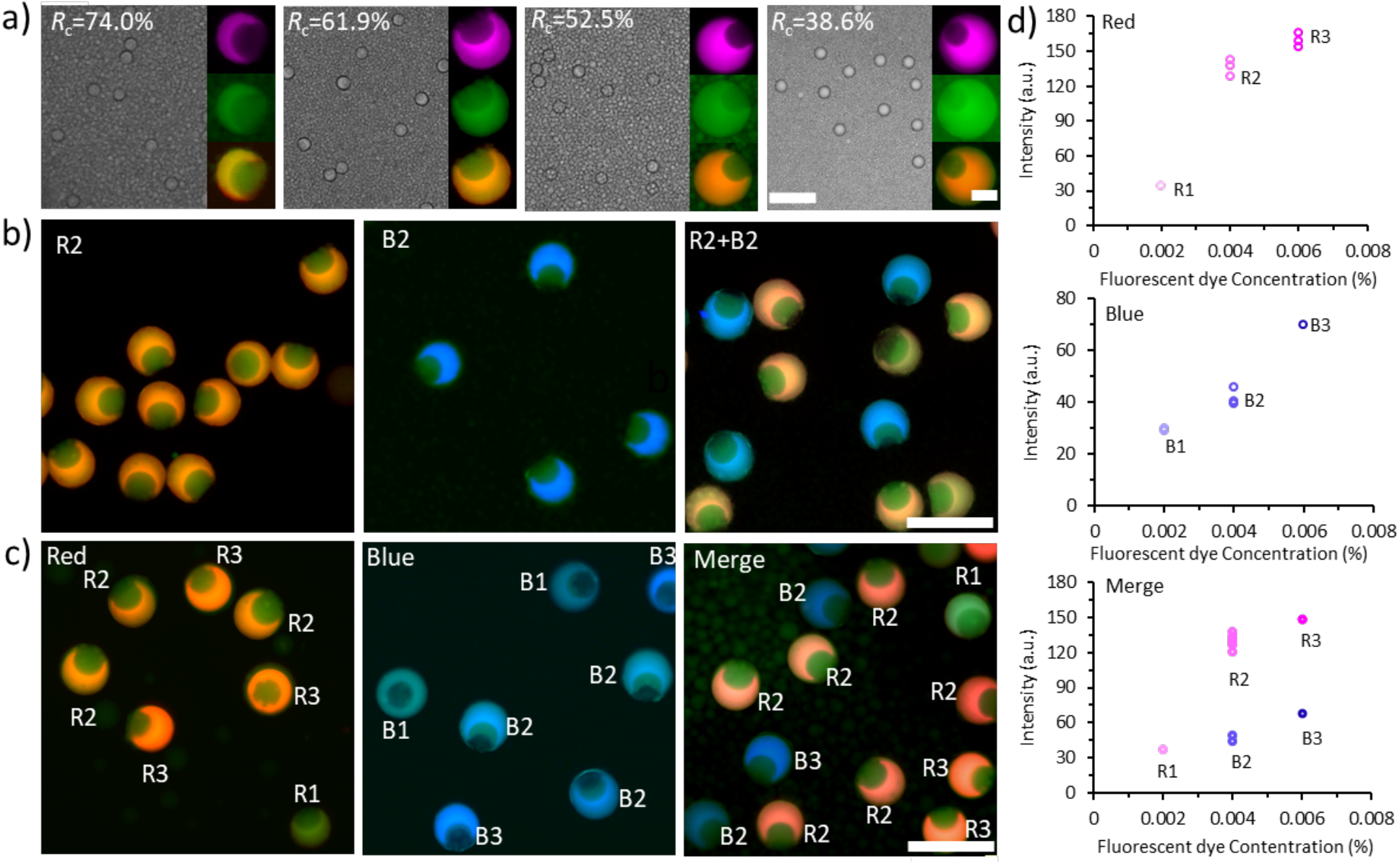
Dropicles formation with fluorescent encoding with variable size crescent particles. (a) Bright-filed (left) and fluorescence (right) microscope images of dropicles formed using fluorescent crescent particles with different *R*_c_ from 38.6% to 74.0%. The scale bars represent 200 μm (left) and 25 μm (right). (b, c) Fluorescence microscope images of dropicles formed using fluorescent crescent particles with different concentrations and types of fluorescent dyes. The scale bars represent 100 μm. The three concentrations (0.002, 0.004, 0.006 w/w%) of red-emitting dye within the crescent particles are labeled as R1, R2, and R3, respectively. The three concentrations (0.002, 0.004, 0.006 w/w%) of blue-emitting dye within the crescent particles are labeled as B1, B2, and B3, respectively. (d) The measured fluorescent intensities for the blue-emitting and red-emitting particles in (c) are plotted with variable concentrations of the dyes.

### 2.6. The effect of pH value on the fluorescent particles

To probe the effect of pH value on fluorescent particles, we incubated them in variable pH bulk solutions and dropicles, and captured the fluorescent images at regular intervals (**Figure 7**a). The fluorescence intensity of blue-emitting and red-emitting particles in pH 1 and pH 7 bulk solutions did not change much within 24 h (Figure 7b,c). The particles that were encapsulated in pH 1 and pH 7 aqueous solution in the form of dropicles also showed a similar stable trend over time. However, the fluorescence intensity of both blue-emitting and red-emitting particles in pH 14 bulk solution decreased dramatically within 1 h (Figure 7d,e and **Figure S7**). The particles were completely disintegrated after 24 h in pH 14 bulk solution; therefore, the fluorescent signal disappeared in the bulk solution due to diffusion. However, the intensity of blue-emitting particles in the pH 14 dropicles only decreased by ∼10% within next 24 h. The blue-emitting particles’ structure within the dropicle remained relatively intact for the first hour, compared to that at pH 14 bulk solution. Since, the dropicles only held ∼1pL volume of the pH 14 aqueous, the degradation of the particle was slower due to the unavailability of fresh high pH solution in abundance.

**Figure 7.**
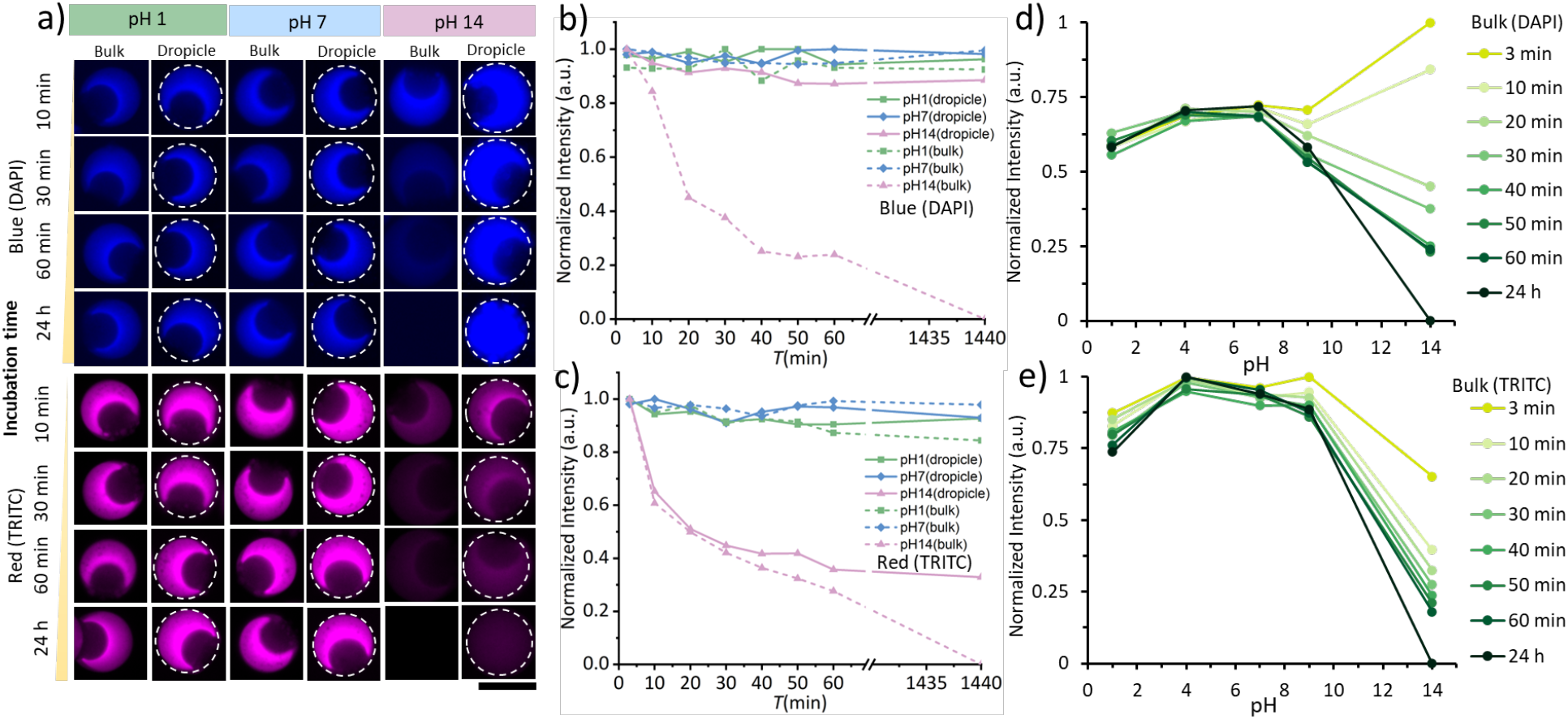
Effect of solution pH on the blue– and red-emitting fluorescent particles. (a) Fluorescent blue and red microscopy images of particles in different pH bulk solutions and dropicles over time. The scale bar represents 50 μm. The normalized intensities of blue-emitting (b) and red-emitting (c) particles in different pH bulk solutions and dropicles (pH 1, 7, 14) as a function of time. (b, c) The fluorescence intensities of samples were normalized against the individual maximum intensity of each sample separately. The normalized intensities of blue-emitting (d) and red-emitting (e) particles in different pH bulk solutions (pH 1, 4, 7, 9, 14) as a function of time. (d, e) The fluorescence intensities of samples were normalized against the overall maximum intensity.

Although the blue-emitting particle structure was also disintegrated after 24 h, a homogenous fluorescence signal was still present within the dropicle. This indicated that the pH 14 solution was not able to quench the blue-emitting fluorescent dye, rather the PEGDA network was completely dismantled. The continuous oil phase contained the degraded particle material precluding the dye from leaking outside of the droplet. These results also demonstrated the strength of the dropicle as a compartmentalization technique retaining an amplified assay signal from tiny volume of reactants and preventing the leakage of analytes. However, the red-emitting particles in the pH 14 dropicles lost the fluorescent signal quite rapidly compared to the blue-emitting particles. The red-emitting rhodamine B fluorophores, once separated from the disintegrating PEGDA network, leaked into the continuous oil phase. The leakage of fluorescent signal from the droplets has been corelated with the surfactant concentration in the oil.^33,54^ A relatively higher leakage rate of the ionic rhodamine B dye is attributed to its interaction with the surfactant at the interface, whereas the neutral blue-emitting dye preferred to remain within the aqueous solution. The red-emitting dye, as long as it was bound to the PEGDA network within pH 1 and pH 7 dropicles, did not lose fluorescent signal, which was consistent with the behavior of blue-emitting particles.

The fluorescence intensity of blue-emitting particles increased with increase in pH value from 1 to 7. The emission spectra of blue-emitting fluorophore that we used red-shifts from blue-to-orange as the pH value was decreased from 7 to 1.^46–48^ Since we have imaged the blue-emitting particles in the same DAPI channel, the fluorescence intensity decreased when the fluorescence spectra of particles in the acidic solution redshifts (i.e. pH decreased from 7 to 1) and *vice versa*. The effect of pH value on the particles that emitted red fluorescence in different pH bulk solutions was also evaluated. The red fluorescence intensity showed an increasing trend as the pH increased from 1 to 4 and decreased as the pH further increased from 4 to 7. The fluorescent particles could tolerate an acidic environment with a pH value as low as 1 for 24 h. The intensity variation with variable pH values in the acid solutions provides the possibility of monitoring the pH change of tiny analytes in dropicles using fluorescent particles as the template. Recording full fluorescence spectra and their analysis should open a more sensitive way for probing the pH within the dropicles.^48^

## 3. Conclusion

In summary, we investigated an aqueous two-phase system comprising of PEGDA (*M*_w_ 700) and dextran (*M*_w_ 40,000) by plotting a comprehensive phase diagram. We fabricated crescent-shaped particles by selectively polymerizing photocurable PEGDA inside a microfluidic droplet generation device. The morphology of the crescent-shaped particles was tailored by tuning the concentrations of polymer precursors, UV intensity, and flow rate ratios. We separately introduced two types of fluorescent dyes into the PEGDA network to enable particle encoding via chemical covalent bonds. The dyes were not washed away as long as the particle hydrogel network was stable. The crescent-shaped particle with variable sizes and fluorescent codes were used as a template to form particle-templated droplets or *dropicles* by a simple pipetting operation. The fluorescent particles demonstrated good stability in a neutral pH or highly acidic solution but completely degraded in a high pH basic solution within 24 h. Therefore, the immobilized encoding dyes within the particle structure were lost in a pH 14 bulk solution. However, the fluorescent blue signal was retained within isolated dropicles even after the particle hydrogel network was completely disintegrated. The fluorescence intensity of blue-emitting particles increased with the increase in the pH value from 1 to 7, indicating a promising tool for pH sensing of miniaturized analytes within pL volume dropicles.

## 4. Experimental section

### 4.1. Materials

Poly(ethylene glycol) diacrylate (PEGDA, Mw=700), dextran (Mw=40,000), lithium pheny1– 2,4,6-trimethylbenzoylphosphinate (LAP), fluorescein isothiocyanate-dextran (FITC-dextran, Mw=40,000), 2% trichloro (1H,1H,2H,2H-perfluorooctyl) silane, perfluoro octanol, fluorescein sodium salt, PluronicF-127, sodium hydroxide solution (c(NaOH)=1, pH 14), hydrogen chloride solution (c(HCl)=1, pH 1), and buffer solution (pH 4, pH 9) were purchased from Sigma-Aldrich. Methacryloxyethyl thiocarbamoyl rhodamine B (MA-RhB) was purchased from Polysciences. Novec 7500 fluorinated oil was purchased from Iolitec. FluoSurf 2w/w% in HFE7500 was purchased from Emulseo. Methacrylate-modified blue-emitting dye, MA-P4VB {2– (methacryloyloxy)propyl 6-(4-methoxy-2,5-bis((E)-2-(pyridin-4-yl)vinyl)phenoxy)hexanoate)}, was synthesized as described in SI (see Figure S8).

### 4.2. Phase separation experiment

To identify the phase separation diagram of PEGDA (Mw 700) and dextran (Mw 40,000), we prepared PEGDA and dextran mixtures of 100 μL volumes using different concentrations of polymer precursors at varied volume ratios, which are listed in Table S1. Each mixture was vortexed to ensure full mixing and centrifuged in a benchtop centrifuge at 6,000 rpm for 30 minutes to induce phase separation. If liquid-liquid phase separation occurred in bulk, a clear line could be observed. The corresponding effective concentrations of PEGDA and dextran were labeled as phase separation points. Finally, the binodal line was plotted based on these points.

### 4.3. Fabrication of microfluidic device

The Y-junction microfluidic device was fabricated using a soft lithography process. Master mold was fabricated on 3-inch silicon wafers by photolithography process. The device height was 80 μm. The width of the nozzle and downstream serpentine channel were 30 μm and 400 μm, respectively (**Figure S9**a). The microfluidic device was replica-molded from the master mold by using the pourable PDMS mixture. The PDMS base and crosslinker were mixed at 10:1 mass ratio, poured onto the silicon wafer, degassed, and cured at 65 °C for 1 h. The PDMS channel and glass slide were cleaned with isopropanol and blow-dried. The PDMS device bonded to a glass slide after oxygen plasma treatment. The boned device was treated with 2% v/v trichloro (1H,1H,2H,2H-perfluorooctyl) silane in Novec 7500 to make channel surfaces hydrophobic. After modification, the device was baked at 75 °C for 1 h to evaporate the residual oil in the channels.

To image particle-templated droplets, a rectangle-shaped PDMS chamber was also fabricated using the same soft lithography method. The height of the chamber was 100 μm. The chamber length and width were 15 mm and 5 mm, respectively (Figure S9b).

### 4.4. Fabrication of crescent-shaped hydrogel particles

Typically, a PEG phase comprised of 40% w/w PEGDA, 1% w/w LAP co-injected into the microfluidic device with a dextran phase comprised of 20% w/w dextran using syringe pumps (Cetoni pump) at a rate of 30-300 μL/h. FITC-dextran was added into the dextran phase to observe the distribution of dextran in droplets. For the fabrication of fluorescent particles, methacrylate-modified blue emitting (MA-P4VB) and red-emitting (MA-RhB) dyes were separately added to PEG phase with three different concentrations (0.002% w/w, 0.004% w/w, 0.006% w/w). An oil phase comprised of Novec 7500 and 0.5% w/w fluorinated surfactant was injected at a rate of 1,000-6,000 μL/h. The PEGDA and dextran solution in water was used as the dispersed aqueous phase, which broke up into monodisperse ATPS droplets by the continuous oil phase. The PEGDA and dextran phases separated on the chip rapidly and formed a core-shell structure. The PEGDA shell was crosslinked on-chip by UV light through a DAPI filter and 5x objective of Leica microscope equipped with a Lumencor Spectra-X light engine (21 mW/mm^2^, 395 nm at 10x magnification). The crosslinked particles were collected after emulsion was broken by 20 % v/v perfluoro-octanol in Novec 7500 at approximately 1:1 volume ratio. A layer of distilled water was added on the top to transfer particles to the aqueous phase. Then excess oil was removed, and samples were washed two times with Novec 7500 to remove the remaining surfactant. In the next step, relatively low-density hexane was added to extract the remaining Novec 7500 oil. The oil layer at the top of the aqueous particle sample was removed by pipetting. Lastly, the particles were washed three times with 70% ethanol and three times with distilled water to remove dextran. The particles were stored in water for later experiments.

### 4.5. Particle-templated droplet formation

The particle-templated droplet or dropicle was formed with a simple workflow based on pipetting. Firstly, the crescent-shaped particles suspended in water were centrifuged at 6,000 rpm for 1 min and the excess water was removed with a micropipette. Then 10 μL of fluorescein sodium aqueous solution was added to the particles and mixed well with vortex (VWR). Next, an oil solution containing Novec 7500 and 0.5% w/w fluorinated surfactant was added to the particle suspensions at about 1:10 volume ratio of aqueous to oil solution. Lastly, the mixture was pipetted using 200 μL micropipette (Eppendorf) for about 3 mins to generate dropicles (∼120 pipetting strokes). The resulting dropicles were transferred to the PDMS chamber, which was filled with Novec 7500 in advance by micropipette for imaging. Moreover, to reduce the loss of particles during the dropicle formation process, all pipette tips and Eppendorf tubes were precoated with 0.05% PluronicF-127 aqueous solution.

### 4.6. Investigation of the effect of pH value on the fluorescent particles

To investigate the effects of pH value on the fluorescent particles in bulk solution, the fluorescent particles suspended in water were transferred to five wells in a well plate, where the medium of each of the four wells was separately exchanged for a different pH solution (pH 1, pH 4, pH 9, pH14) after three consecutive washes. The fluorescent particles were in the water as a control group (pH 7). The same protocol was used for two types of fluorescent particles that emitted blue and red fluorescence. For the investigation of the effects of pH value on the fluorescent particles in dropicle, the blue– and red-emitting fluorescent particles were separately mixed with different pH aqueous solutions (pH 1, pH 7, pH 14) using the dropicle formation protocol. Novec 7500 oil with 0.5% w/w fluorinated surfactant was added and the mixtures were pipetted about 120 times with a 200 μL micropipette to create dropicles. The fluorescence intensity of particles in bulk solutions and dropicles was collected every 10 min in 1 h. After 24 h incubation time at room temperature, the fluorescence intensity was measured again.

### 4.7. Swelling property experiment

The crescent-shaped particle suspension was placed on glass slides to dry at room temperature for 50 h. Following the water was added to dried particles to rehydrate it. The dried and swollen particle images were captured by microscope to analyze the size change of particles during the rehydration process.

### 4.8. Characterization

The morphologies of droplets, crescent-shaped particles, and dropicles were characterized by a microscope (Thunder imager, Leica Microsystem) and a Scanning Electron Microscope (SEM) (JEOL). The size distribution of droplets, particles, and dropicles was analyzed using MATLAB and Image J. The fluorescent particles with blue-emitting dye were imaged in the DAPI channel (excitation (ex) filter 395nm / dichroic mirror (dm) 415nm / emission (em) filter 430nm), and the particles with red-emitting dye were imaged in the TRITC channel (ex. filter 555nm / dm 570nm / em. filter 595nm). The fluorescein-loaded dropicles were imaged in the FITC channel (ex. filter 475nm / dm 490nm / em. filter 515nm).

### 4.9 Synthesis of MA-P4VB dye

The parent chromophore, sodium 6-(4-methoxy-2,5-bis((E)-2-(pyridin-4-yl)vinyl)phenoxy) hexanoate (c-P4VB), was prepared according to the previously published procedure.^47^ It was further converted into N-hydroxysuccinimide-derivative (NHS-P4VB) and subsequently into MA-derivative (MA-P4VB) as follows (**Figure S10**).

*Synthesis of 2,5-dioxopyrrolidin-1-yl 6-(4-methoxy-2,5-bis((E)-2-(pyridin-4-yl)vinyl)phenoxy) hexanoate (NHS-P4VB):*

c-P4VB was reacted with N-hydroxysuccinimide in presence of a carbodiimide reagent EDC (1– ethyl-3-(3-dimethylaminopropyl)carbodiimide hydrochloride) and triethylamine in dry THF. For this, 200 mg (0.43 mmol) of c-P4VB, 170 mg (0.88 mmol) EDC and 91 mg (0.90 mmol) triethylamine were mixed in 10 ml of anhydrous tetrahydrofuran and stirred vigorously overnight, resulting in a fine yellow suspension. This was taken into a gas-tight glass syringe and filtered through a PTFE syringe filter (0.45 µm). The filtrate was evaporated to dryness, the resulting solid was washed quickly but thoroughly with water and collected by suction filtration. After drying under vacuum, 198 mg (85 %) of yellow-orangeish NHS-P4VB was obtained.

^1^H NMR (Bruker Avance Neo 400 MHz, CDCl_3_), δ (ppm): 8.60-8.55 (m, 4H), 7.67 (d, ^3^*J* = 16.5 Hz, 1H), 7.66 (d, ^3^*J* = 16.5 Hz, 1H), 7.41-7.37 (m, 4H), 7.13 (2s, 2H), 7.06 (d, ^3^*J* = 16.5 Hz, 1H), 7.05 (d, ^3^*J* = 16.5 Hz, 1H), 4.09 (t, ^3^*J* = 6.3 Hz, 2H), 3.94 (s, 3H), 2.79 (s, 4H, succinimide-moiety) 2.68 (t, ^3^*J* = 7.2 Hz, 2H), 1.98-1.83 (m, 4H), 1.74-1.64 (m, 2H).

ESI-MS (acetonitrile): m/z = 542.5 (MH^+^, 100%)

*Synthesis of 2-(methacryloyloxy)propyl 6-(4-methoxy-2,5-bis((E)-2-(pyridin-4-yl)vinyl)phenoxy) hexanoate (MA-P4VB):*

In a 20 ml vial equipped with a septum and a stirring bar, 15 mg DMAP (N,N-dimethylamino-4-pyridine, 0.12 mmol), 130 mg NHS-P4VB (0.24 mmol) and 3 ml HPMA (2-hydroxypropyl/1-hydroxyisopropyl methacrylate, mixture of isomers, 97%, stabilized, 144 g/mol, 266 mg, 22 mmol) were dissolved in 3 ml of anhydrous THF. The formed suspension was stirred tightly closed and protected from light for 7 days at room temperature. The reaction was stopped by removal of THF under vacuum followed by a transfer of the mixture into 35 ml water in a centrifugation vessel. 5 ml acetone were additionally used to transfer the complete reaction mixture into the centrifugation vial. After centrifugation, the viscous honey-like residue was re-dissolved in 5 ml acetone and precipitated by addition of 35 ml water. This was centrifuged, the water was discarded and the residue transferred into a glass vial by means of dissolving with acetone. The acetone was removed in vacuo leaving 125 mg (91% yield) of solid material. The product is an approx. 2:1 mixture of two isomers due to the methacryloyloxypropyl-group, with the major isomer shown in Figure S10.

^2^H NMR (Bruker Avance Neo 400 MHz, CDCl_3_), δ (ppm): 8.63-8.57 (m, 4H), 7.69 (d, ^3^*J* = 16.4 Hz, 2H), 7.45-7.38 (m, 4H), 7.18-7.03 (m, 4H), 6.11 (m, 1H), 5.57 (m, 1H), 5.26-5.17 (m, 1H), 4.27-4.05 (m, 4H), 3.97 (s, 3H), 2.44-2.36 (m, 2H), 2.00-1.54 (m, 9H) 1.32-1.25 (m, 3H).

ESI-MS (acetonitrile): m/z = 570 (M^+^, 100%)

See **Figure S11** for ESI-MS spectrum of MA-P4VB and Figure **S12** for NMR (CDCl_3_) spectrum of MA-P4VB.

## Acknowledgments

Y.Y. acknowledges the Chinese Scholarship Counsel for the doctoral fellowship. The authors acknowledge Mr. Furkan Ginaz Almus for support in data analysis. Finally, the authors would like to acknowledge the support of all staff members and researchers working in the facilities of the Heinz Nixdorf Chair of Biomedical Electronics and the Central Electronics and Information Technology Laboratory – ZEIT^lab^.

## Authors contributions

Y.Y. and G.D. conceptualized the work. Y.Y. performed the experiments, analyzed the results, and wrote the first draft of the manuscript. S.I.V., and B.R. assisted in the experiments and data analysis. Y.Y., S.I.V., B.R., and G.D. revised the manuscript. G.D. supervised the work.

## Supporting Information

**Table S1.** The concentrations of PEGDA 700 and dextran 40,000 for the phase separation experiment in bulk.

| Initial PEGDA concentration | Initial dextran concentration | Initial dextran volume (μL) | Initial PEGDA volume (μL) | Effective dextran concentration | Effective PEGDA concentration | phase separation |
| --- | --- | --- | --- | --- | --- | --- |
| 0.5 | 0.4 | 50 | 50 | 0.2 | 0.25 | Y |
| 0.5 | 0.4 | 40 | 60 | 0.16 | 0.3 | Y |
| 0.5 | 0.4 | 30 | 70 | 0.12 | 0.35 | Y |
| 0.5 | 0.4 | 20 | 80 | 0.08 | 0.4 | Y |
| 0.5 | 0.3 | 70 | 30 | 0.21 | 0.15 | Y |
| 0.5 | 0.3 | 60 | 40 | 0.18 | 0.2 | Y |
| 0.5 | 0.3 | 50 | 50 | 0.15 | 0.25 | Y |
| 0.5 | 0.3 | 40 | 60 | 0.12 | 0.3 | Y |
| 0.5 | 0.3 | 30 | 70 | 0.09 | 0.35 | Y |
| 0.5 | 0.3 | 20 | 80 | 0.06 | 0.4 | Y |
| 0.5 | 0.3 | 10 | 90 | 0.03 | 0.45 | Y |
| 0.5 | 0.2 | 70 | 30 | 0.14 | 0.15 | Y |
| 0.5 | 0.2 | 60 | 40 | 0.12 | 0.2 | Y |
| 0.5 | 0.2 | 50 | 50 | 0.1 | 0.25 | Y |
| 0.5 | 0.2 | 40 | 60 | 0.08 | 0.3 | Y |
| 0.5 | 0.2 | 30 | 70 | 0.06 | 0.35 | Y |
| 0.5 | 0.2 | 20 | 80 | 0.04 | 0.4 | Y |
| 0.5 | 0.15 | 60 | 40 | 0.09 | 0.2 | Y |
| 0.5 | 0.15 | 70 | 30 | 0.105 | 0.15 | Y |
| 0.5 | 0.15 | 65 | 35 | 0.0975 | 0.175 | Y |
| 0.5 | 0.1 | 70 | 30 | 0.07 | 0.15 | N |
| 0.5 | 0.1 | 65 | 35 | 0.065 | 0.175 | Y |
| 0.5 | 0.1 | 60 | 40 | 0.06 | 0.2 | Y |
| 0.5 | 0.1 | 50 | 50 | 0.05 | 0.25 | Y |
| 0.5 | 0.1 | 40 | 60 | 0.04 | 0.3 | Y |
| 0.5 | 0.1 | 30 | 70 | 0.03 | 0.35 | Y |
| 0.5 | 0.05 | 55 | 45 | 0.0275 | 0.225 | Y |
| 0.5 | 0.05 | 60 | 40 | 0.03 | 0.2 | Y |
| 0.5 | 0.035 | 45 | 55 | 0.01575 | 0.275 | Y |
| 0.5 | 0.035 | 55 | 45 | 0.01925 | 0.225 | Y |
| 0.4 | 0.2 | 70 | 30 | 0.14 | 0.12 | N |
| 0.25 | 0.4 | 20 | 80 | 0.08 | 0.2 | Y |
| 0.25 | 0.4 | 30 | 70 | 0.12 | 0.175 | Y |
| 0.25 | 0.4 | 40 | 60 | 0.16 | 0.15 | Y |
| 0.25 | 0.4 | 50 | 50 | 0.2 | 0.125 | Y |
| 0.25 | 0.4 | 60 | 40 | 0.24 | 0.1 | Y |
| 0.25 | 0.2 | 70 | 30 | 0.14 | 0.075 | N |
| 0.25 | 0.2 | 60 | 40 | 0.12 | 0.1 | N |
| 0.25 | 0.2 | 50 | 50 | 0.1 | 0.125 | N |
| 0.25 | 0.2 | 40 | 60 | 0.08 | 0.15 | N |
| 0.25 | 0.2 | 30 | 70 | 0.06 | 0.175 | Y |
| 0.25 | 0.2 | 20 | 80 | 0.04 | 0.2 | Y |
| 0.25 | 0.2 | 25 | 75 | 0.05 | 0.1875 | Y |
| 0.25 | 0.2 | 35 | 65 | 0.07 | 0.1625 | Y |
| 0.25 | 0.1 | 70 | 30 | 0.07 | 0.075 | N |
| 0.25 | 0.1 | 60 | 40 | 0.06 | 0.1 | N |
| 0.25 | 0.1 | 50 | 50 | 0.05 | 0.125 | N |
| 0.25 | 0.1 | 40 | 60 | 0.04 | 0.15 | N |
| 0.25 | 0.1 | 30 | 70 | 0.03 | 0.175 | N |
| 0.25 | 0.1 | 20 | 80 | 0.02 | 0.2 | Y |
| 0.25 | 0.05 | 70 | 30 | 0.035 | 0.075 | N |
| 0.25 | 0.05 | 60 | 40 | 0.03 | 0.1 | N |
| 0.25 | 0.05 | 50 | 50 | 0.025 | 0.125 | N |
| 0.25 | 0.05 | 40 | 60 | 0.02 | 0.15 | N |
| 0.25 | 0.05 | 30 | 70 | 0.015 | 0.175 | N |
| 0.25 | 0.05 | 20 | 80 | 0.01 | 0.2 | Y |
| 0.25 | 0.05 | 10 | 90 | 0.005 | 0.225 | Y |

**Figure S1.**
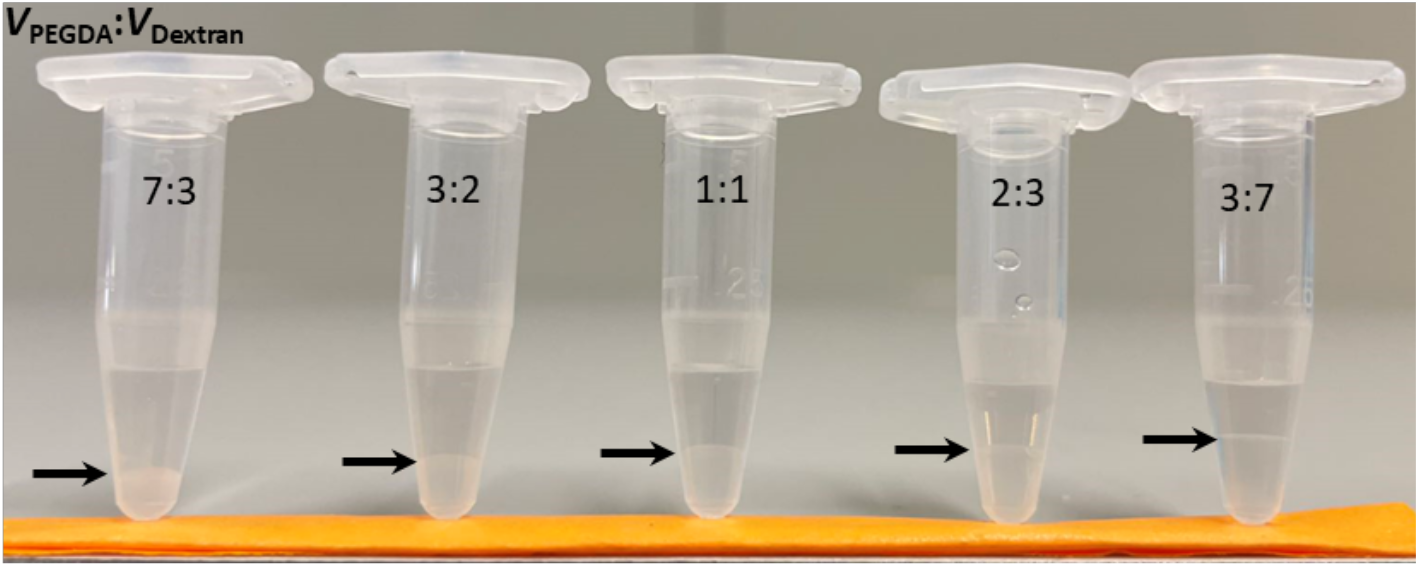
Demonstration of phase separation in bulk samples. The samples were prepared from a mixture of 30% w/w dextran and 50% w/w PEGDA at different volume ratios.

**Figure S2.**
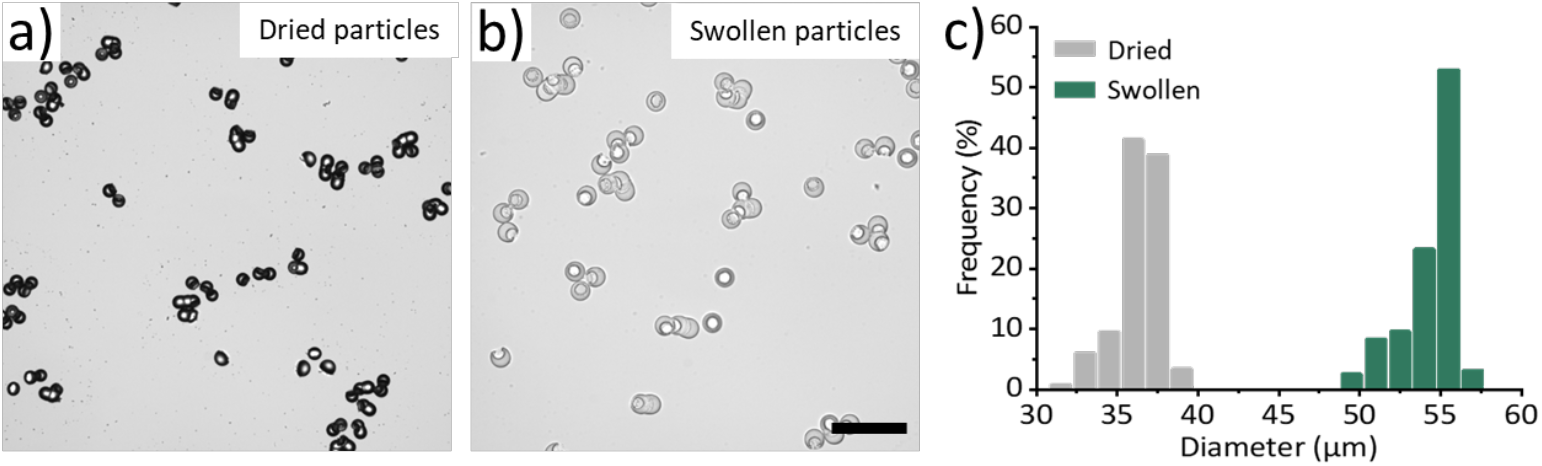
Microscope images of (a) dried particles and (b) swollen particles after rehydration. The scale bar represents 200 μm. (c) The histogram of dried particles and swollen particles.

**Figure S3.**
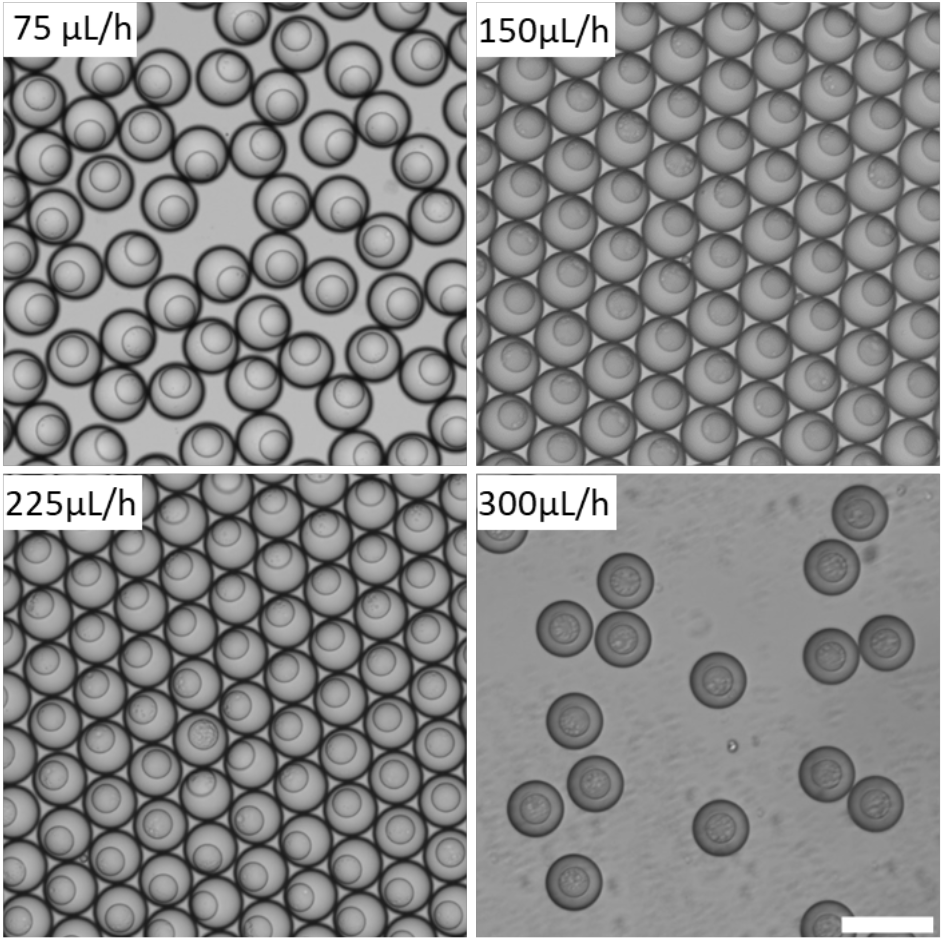
Microscope images of droplets generated at four different aqueous flow rates (75-300 μL/h). The scale bar represents 200 μm.

**Figure S4.**
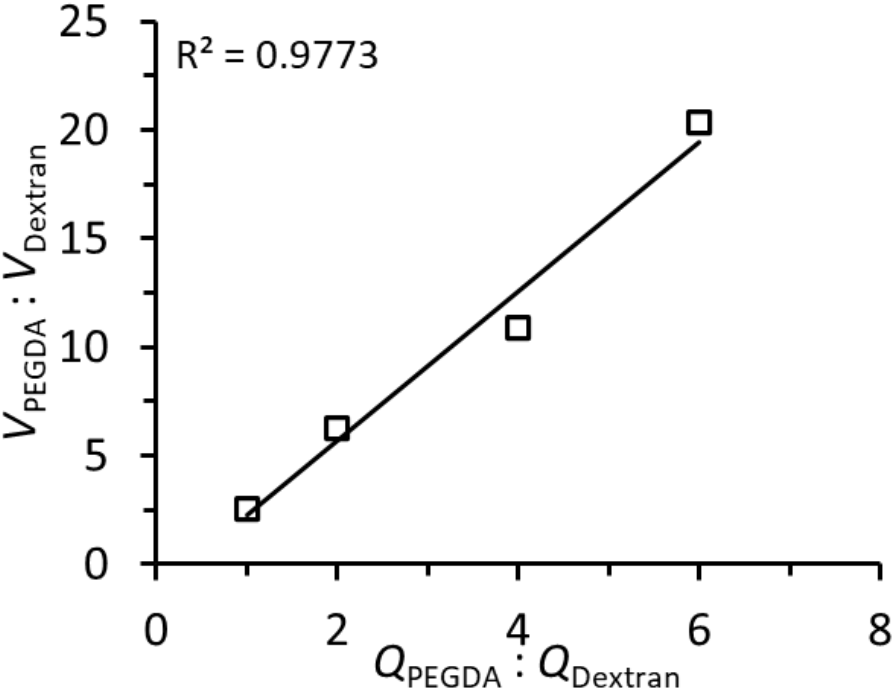
The volume ratio of PEGDA and dextran (*V*_PEGDA_:*V*_Dextran_) of the droplet as a function of the flow rate ratio of PEGDA and dextran.

**Figure S5.**
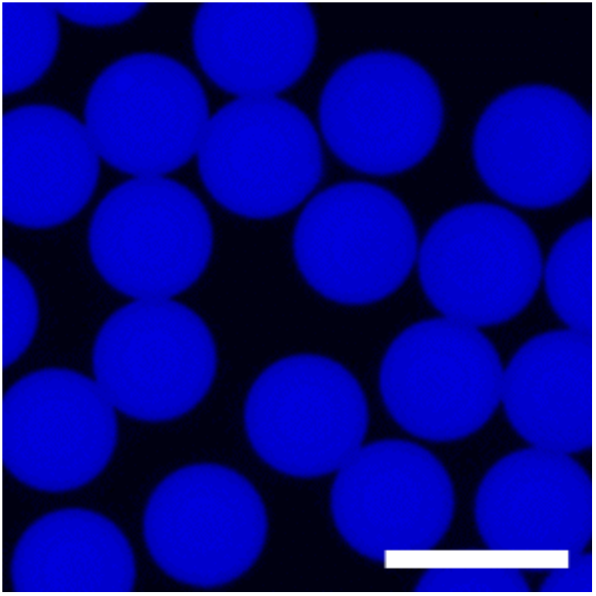
Fluorescent microscope image of blue-emitting dye distribution in aqueous droplets. The scale bar is 100 μm.

**Figure S6.**
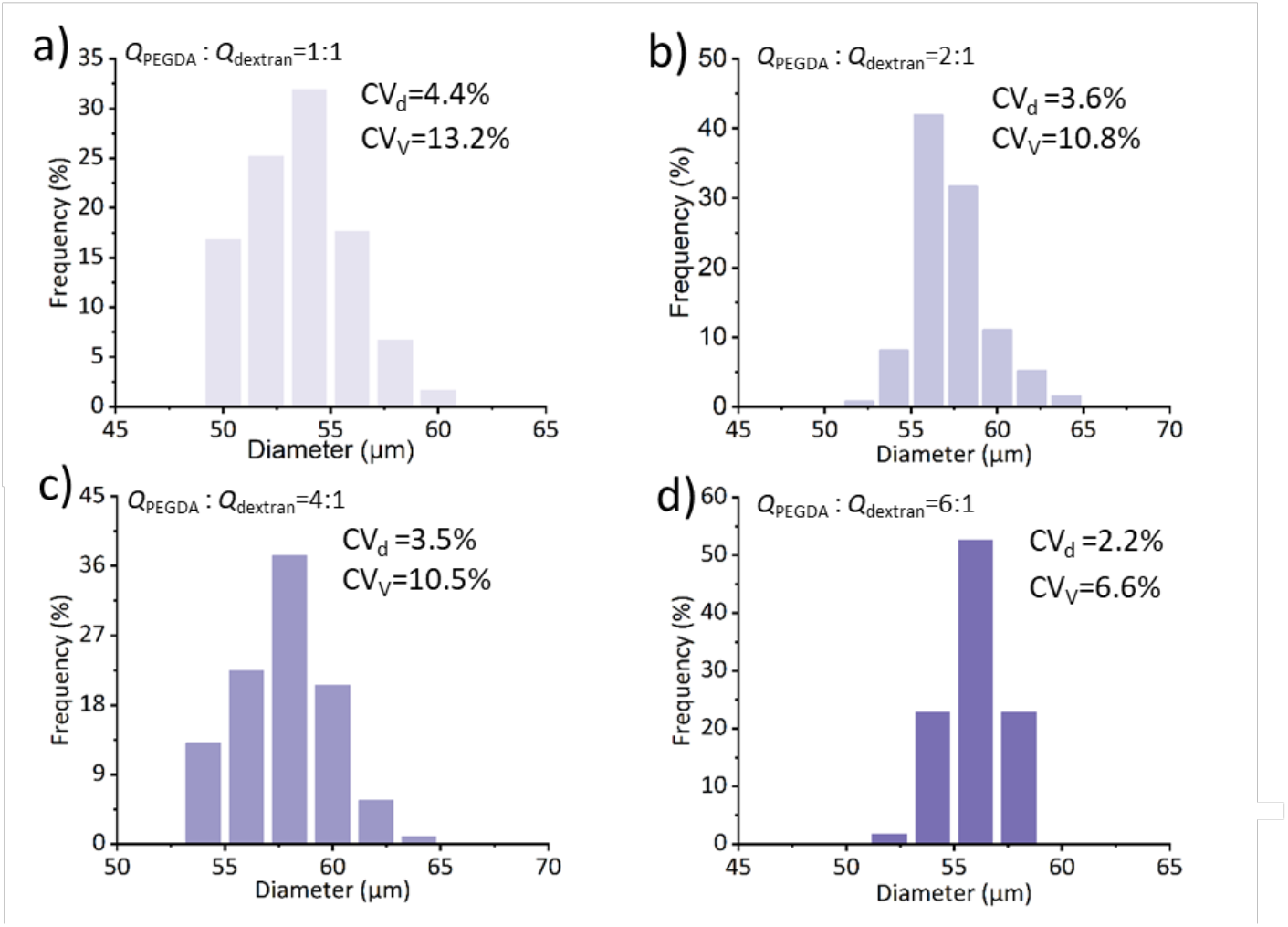
(a-d) Histograms of dropicles formed using fluorescent crescent particles fabricated with different *Q*_PEGDA_: *Q*_Dextran_ from 1:1 to 6:1.

**Figure S7.**
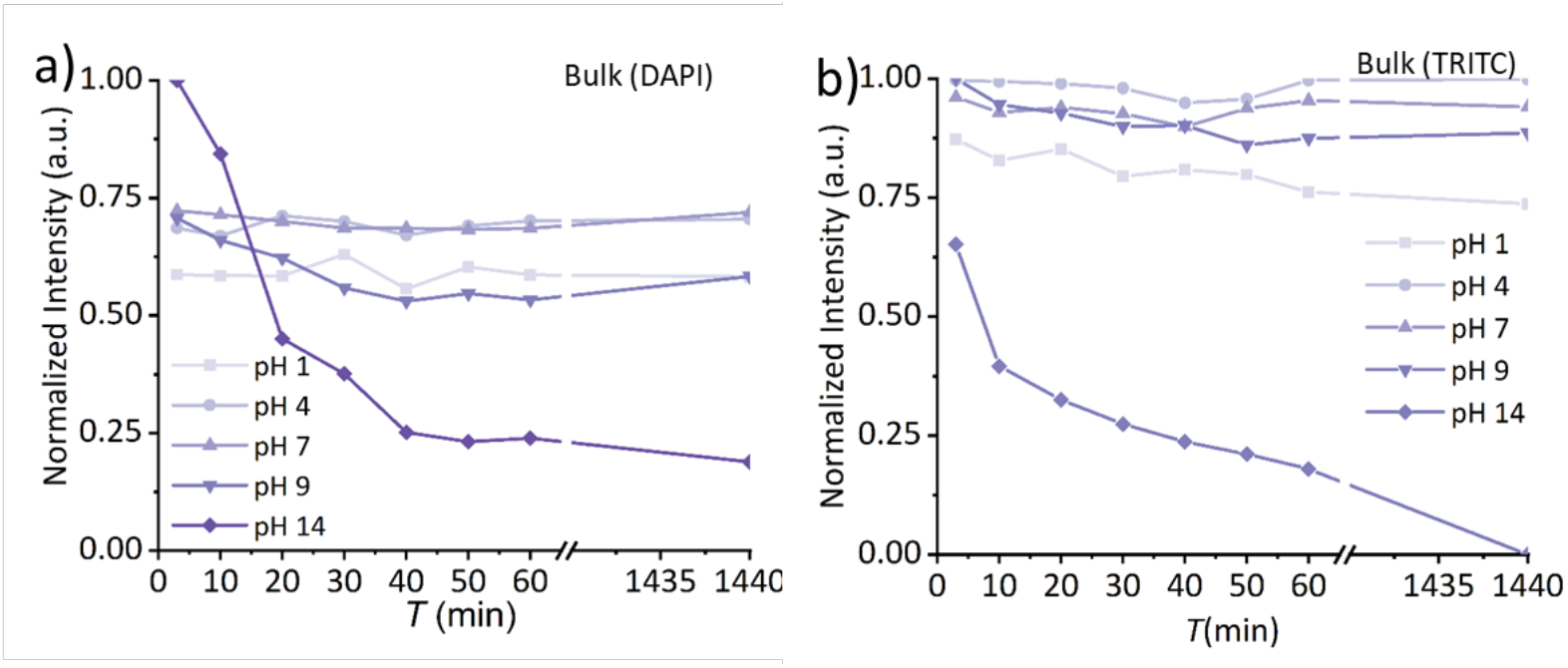
Effect of pH value on the blue-emitting and red-emitting fluorescent particles. The normalized intensity of the blue-emitting (a) and red-emitting (b) fluorescent particles in different pH bulk solutions as a function of time. (a, b)The fluorescence intensities of samples were normalized against the overall maximum intensity.

**Figure S8.**
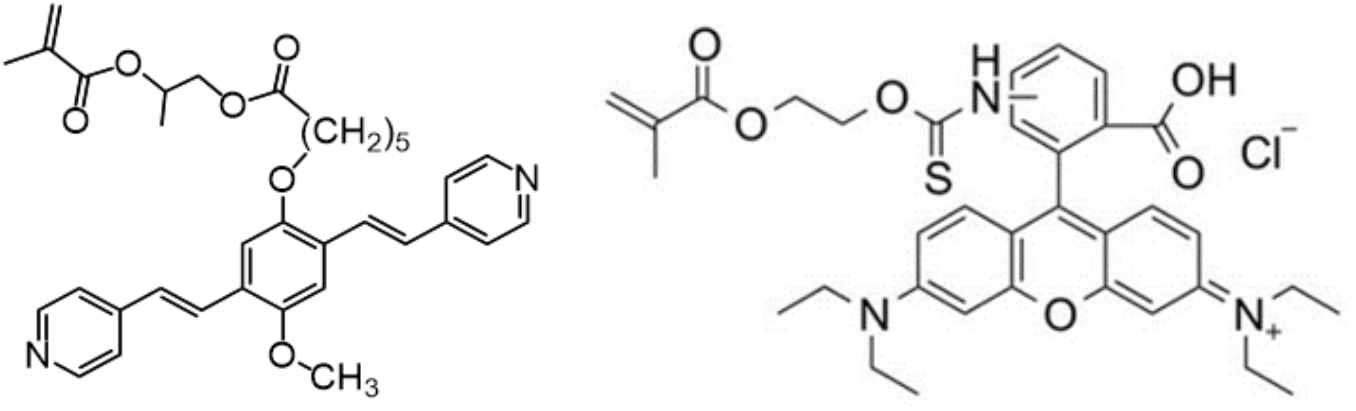
Chemical structures of blue-emitting (left) and red-emitting (right) dyes used for copolymerization with PEGDA in the particles.

**Figure S9.**
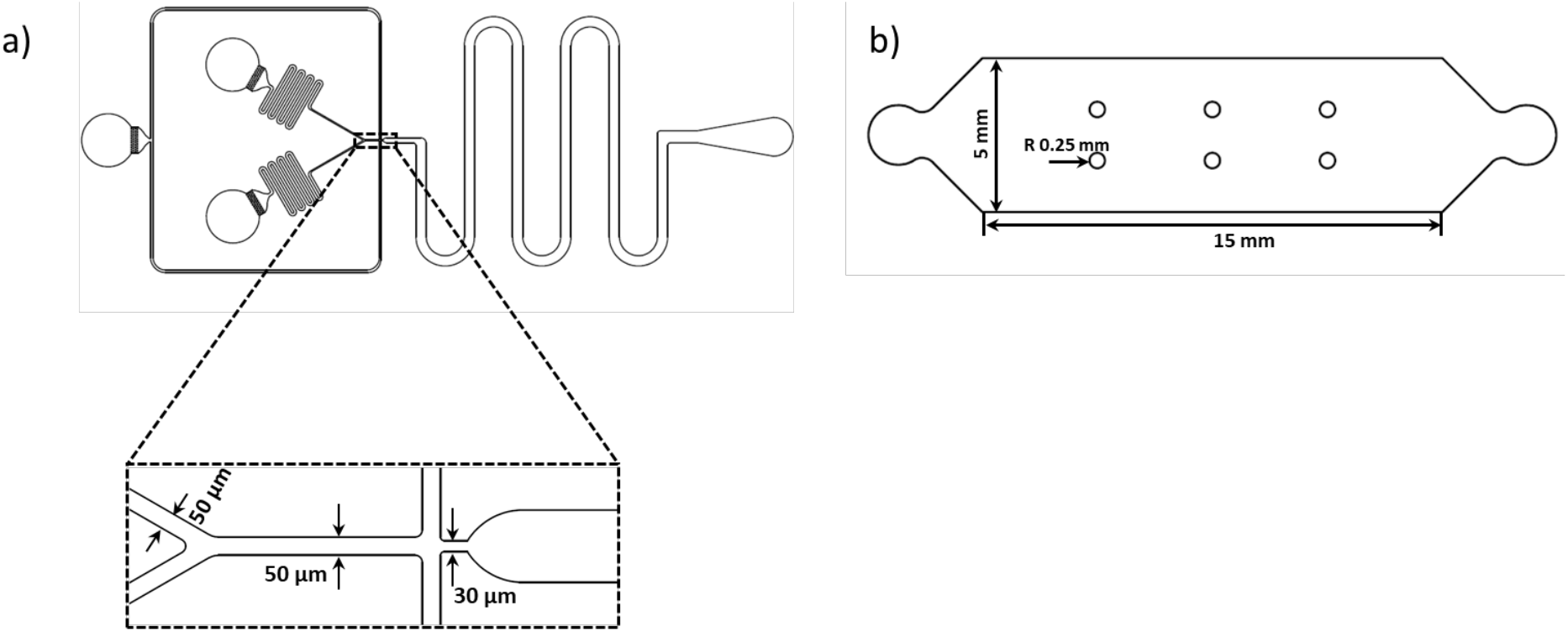
The CAD layouts of the droplet generation device (a) and observation chamber (b).

**Figure S10.**
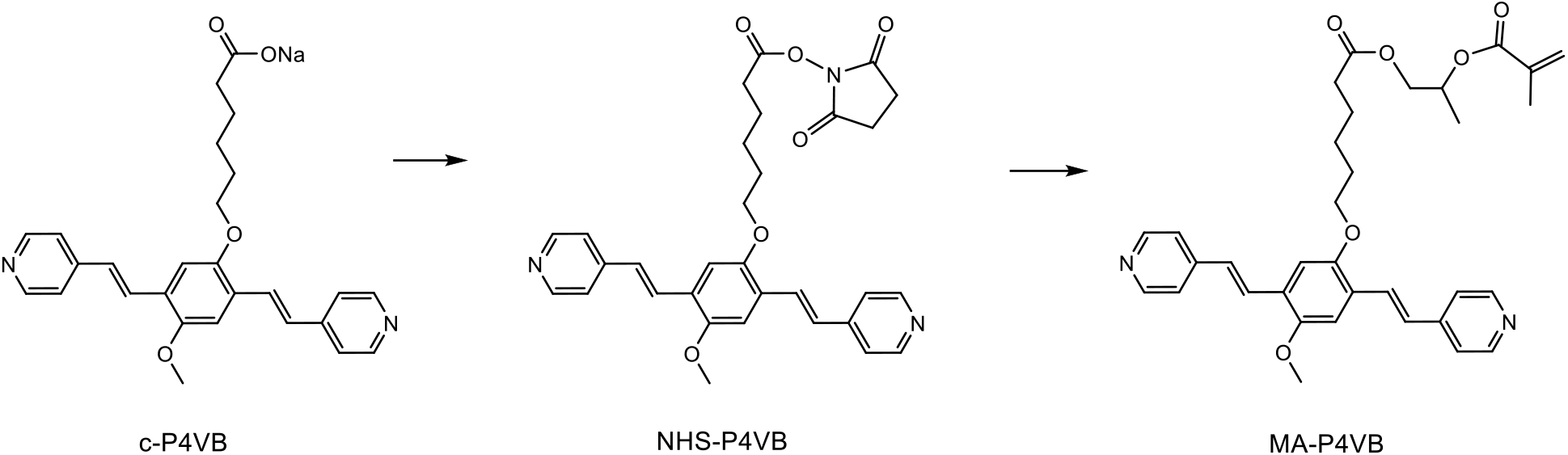
Conversion of c-P4VB to NHS-P4VB to MA-P4VB.

**Figure S11:**
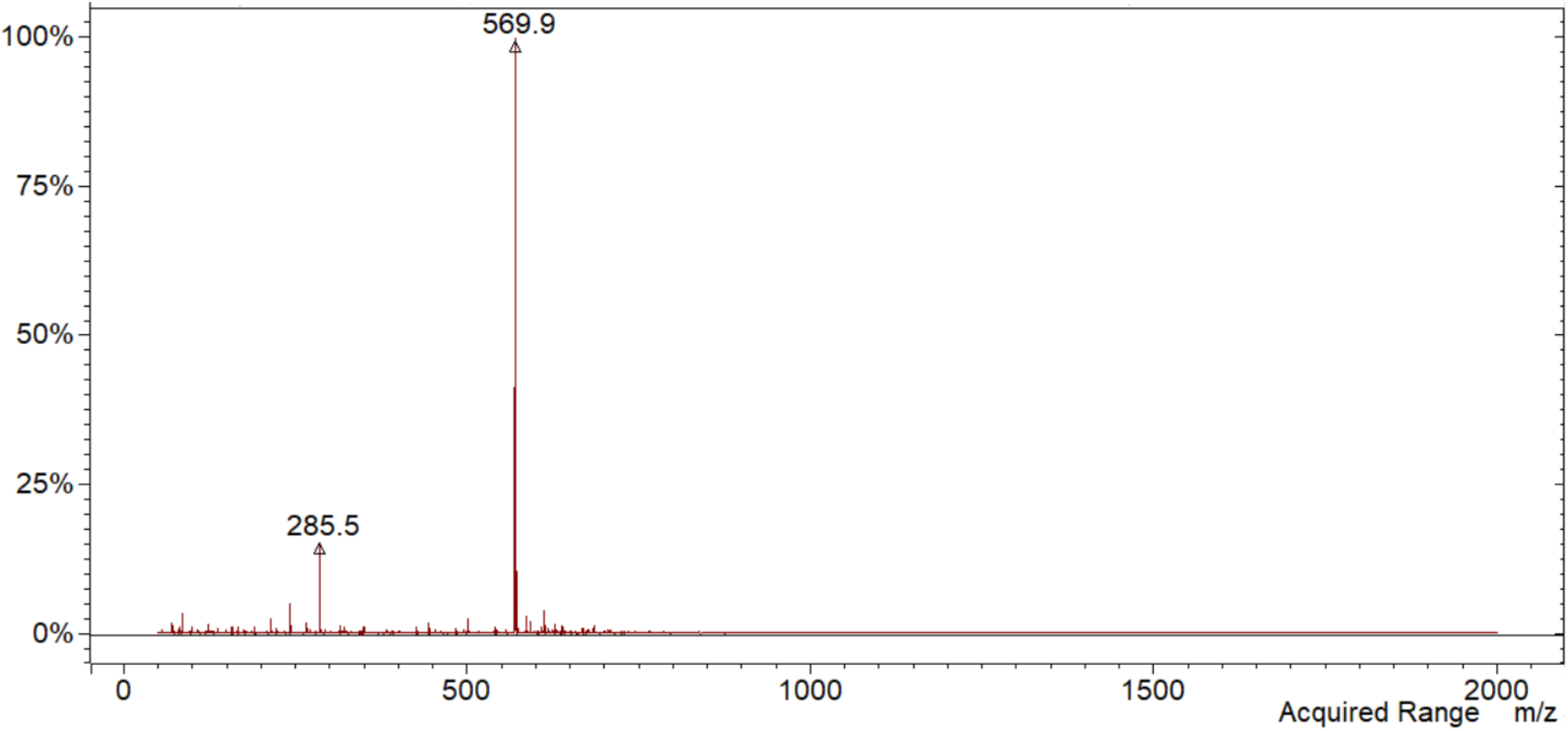
ESI-MS spectrum of MA-P4VB recorded in acetonitrile and showing the molecular peak of MA-P4VB (m/z = 570). The peak at 285 m/z is an adduct of DMAP with hydroxypropyl methacrylate, which is over-represented. No signals of this adduct could be observed by ^1^H NMR spectroscopy.

**Figure S12:**
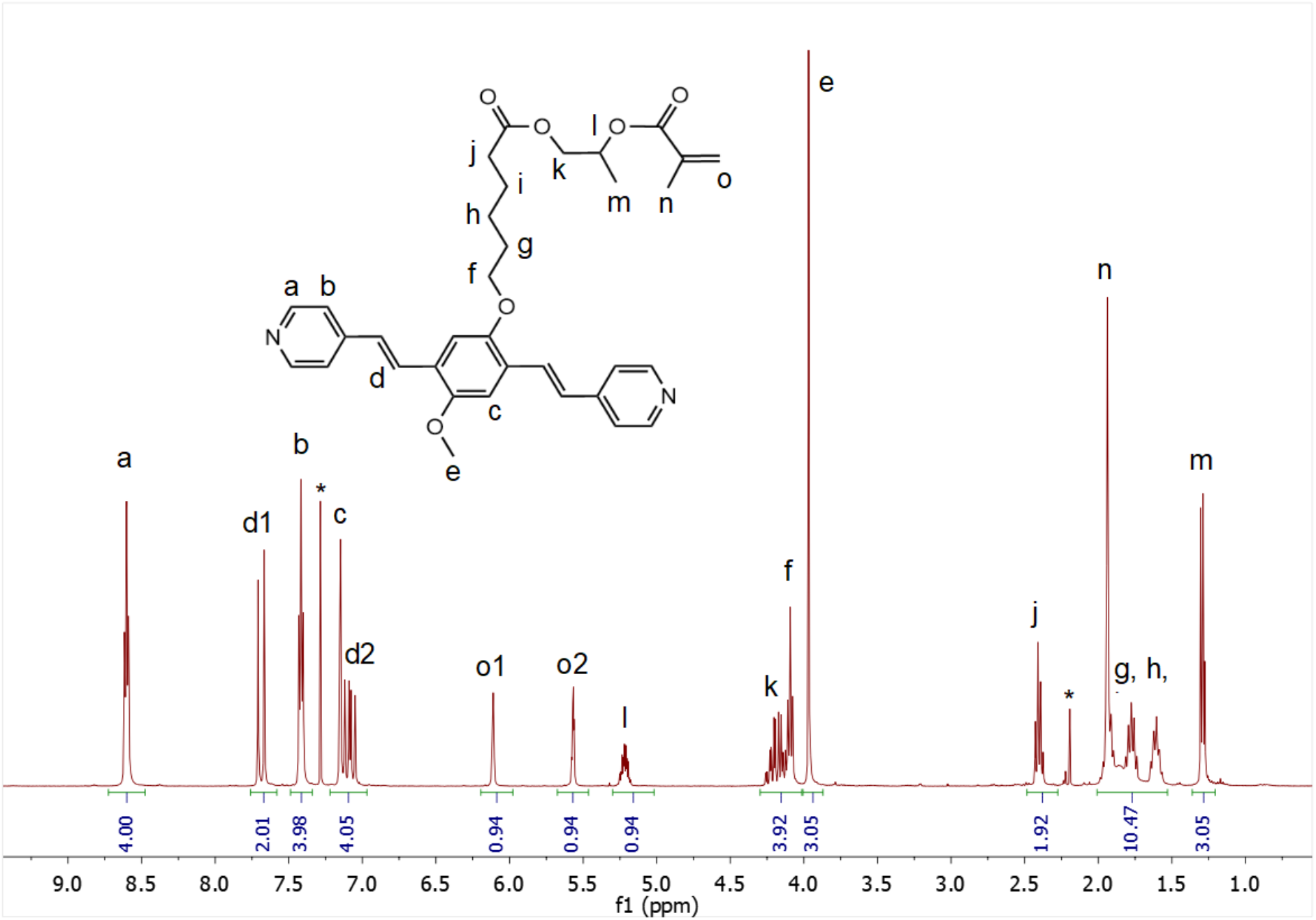
^2^H NMR (CDCl_3_) spectrum of MA-P4VB with the assignment of the signals. Peaks from solvent impurities (CHCl_3_, acetone) are marked with asterisk. The integral 10.47 is overestimated (expected 9H) due to water protons underneath, coming from the deuterated solvent.

## Reference

1. Wu, C., Dougan, T. J. & Walt, D. R. High-Throughput, High-Multiplex Digital Protein Detection with AAomolar Sensitivity. ACS Nano 16, 1025–1035 (2022).

2. Fan, R. et al. Integrated barcode chips for rapid, multiplexed analysis of proteins in microliter quantities of blood. Nat Biotechnol 26, 1373–1378 (2008).

3. Rissin, D. M. et al. Single-molecule enzyme-linked immunosorbent assay detects serum proteins at subfemtomolar concentrations. Nat Biotechnol 28, 595–599 (2010).

4. Shim, J. U. et al. Ultrarapid generation of femtoliter microfluidic droplets for single-molecule-counting immunoassays. ACS Nano 7, 5955–5964 (2013).

5. Novak, R. et al. Single-cell multiplex gene detection and sequencing with microfluidically generated agarose emulsions. Angewandte Chemie – International Edition 50, 390–395 (2011).

6. Edd, J. F. et al. Controlled encapsulation of single-cells into monodisperse picolitre drops. Lab Chip 8, 1262–1264 (2008).

7. Mazutis, L., et al. Single-cell analysis and sorting using droplet-based microfluidics. Nat Protoc 8, 870–891 (2013).

8. Gérard, A. et al. High-throughput single-cell activity-based screening and sequencing of antibodies using droplet microfluidics. Nat Biotechnol 38, 715–721 (2020).

9. Jin, A. et al. A rapid and effcient single-cell manipulation method for screening antigen-specific antibody-secreting cells from human peripheral blood. Nat Med 15, 1088–1092 (2009).

10. Zhu, Q. et al. Digital PCR on an integrated self-priming compartmentalization chip. Lab Chip 14, 1176–1185 (2014).

11. Park, H., Kim, H. & Doh, J. Multifunctional Microwell Arrays for Single Cell Level Functional Analysis of Lymphocytes. Bioconjug Chem 29, 672–679 (2018).

12. Joo, B., Hur, J., Kim, G. B., Yun, S. G. & Chung, A. J. Highly Effcient Transfection of Human Primary T Lymphocytes Using Droplet-Enabled Mechanoporation. ACS Nano 15, 12888–12898 (2021).

13. Sesen, M., Alan, T. & Neild, A. Droplet control technologies for microfluidic high throughput screening (μHTS). Lab on a Chip vol. 17 2372–2394 Preprint at 10.1039/c7lc00005g (2017).

14. Brenker, J. C., Collins, D. J., Van Phan, H., Alan, T. & Neild, A. On-chip droplet production regimes using surface acoustic waves. Lab Chip 16, 1675–1683 (2016).

15. Guo, M. T., Rotem, A., Heyman, J. A. & Weitz, D. A. Droplet microfluidics for high-throughput biological assays. Lab on a Chip vol. 12 2146–2155 Preprint at 10.1039/c2lc21147e (2012).

16. Matuła, K., Rivello, F. & Huck, W. T. S. Single-Cell Analysis Using Droplet Microfluidics. Advanced Biosystems vol. 4 Preprint at 10.1002/adbi.201900188 (2020).

17. Mongersun, A., Smeenk, I., Pratx, G., Asuri, P. & Abbyad, P. Droplet Microfluidic Plaoorm for the Determination of Single-Cell Lactate Release. Anal Chem 88, 3257–3263 (2016).

18. Ding, Y., Choo, J. & deMello, A. J. From single-molecule detection to next-generation sequencingti microfluidic droplets for high-throughput nucleic acid analysis. Microfluid Nanofluidics 21, (2017).

19. Ko, J., et al. Single Extracellular Vesicle Protein Analysis Using Immuno-Droplet Digital Polymerase Chain Reaction Amplification. Adv Biosyst 4, (2020).

20. Sjostrom, S. L. et al. High-throughput screening for industrial enzyme production hosts by droplet microfluidics. Lab Chip 14, 806–813 (2014).

21. Stucki, A., Vallapurackal, J., Ward, T. R. & DiArich, P. S. Droplet Microfluidics and Directed Evolution of Enzymes: An Intertwined Journey. Angewandte Chemie – International Edition vol. 60 24368– 24387 Preprint at 10.1002/anie.202016154 (2021).

22. Dittrich, P. S. & Manz, A. Lab-on-a-chip: Microfluidics in drug discovery. Nature Reviews Drug Discovery vol. 5 210–218 Preprint at 10.1038/nrd1985 (2006).

23. D’Amico, C. I., Polasky, D. A., Steyer, D. J., Ruotolo, B. T. & Kennedy, R. T. Ion Mobility-Mass Spectrometry Coupled to Droplet Microfluidics for Rapid Protein Structure Analysis and Drug Discovery. Anal Chem 94, 13084–13091 (2022).

24. Dimatteo, R. & Di Carlo, D. IL-2 secretion-based sorting of single T cells using high-throughput microfluidic on-cell cytokine capture. Lab Chip 22, 1576–1583 (2022).

25. Delley, C. L. & Abate, A. R. Modular barcode beads for microfluidic single cell genomics. Sci Rep 11, (2021).

26. Gu, Z. et al. Bead-Based Multiplexed Droplet Digital Polymerase Chain Reaction in a Single Tube Using Universal Sequences: An Ultrasensitive, Cross-Reaction-Free, and High-Throughput Strategy. ACS Sens 7, 2759–2766 (2022).

27. Zilionis, R. et al. Single-cell barcoding and sequencing using droplet microfluidics. Nat Protoc 12, 44–73 (2017).

28. Collins, D. J., Neild, A., deMello, A., Liu, A. Q. & Ai, Y. The Poisson distribution and beyond: Methods for microfluidic droplet production and single cell encapsulation. Lab on a Chip vol. 15 3439–3459 Preprint at 10.1039/c5lc00614g (2015).

29. De Rutte, J. et al. Suspendable Hydrogel Nanovials for Massively Parallel Single-Cell Functional Analysis and Sorting. ACS Nano (2021) doi:10.1021/acsnano.1c11420.

30. Destgeer, G., Ouyang, M., Wu, C. Y. & Di Carlo, D. Fabrication of 3D concentric amphiphilic microparticles to form uniform nanoliter reaction volumes for amplified affnity assays. Lab Chip 20, 3503–3514 (2020).

31. Wu, C.-Y., et al. Monodisperse drops templated by 3D-structured microparticles. Sci. Adv vol. 6 https://www.science.org (2020).

32. Hatori, M. N., Kim, S. C. & Abate, A. R. Particle-Templated Emulsification for Microfluidics-Free Digital Biology. Anal Chem 90, 9813–9820 (2018).

33. Wang, Y. et al. Counting of enzymatically amplified affnity reactions in hydrogel particle-templated drops. Lab Chip 21, 3438–3448 (2021).

34. Clark, I. C. et al. Microfluidics-free single-cell genomics with templated emulsification. Nat Biotechnol (2023) doi:10.1038/s41587-023-01685-z.

35. Destgeer, G., Ouyang, M. & Di Carlo, D. Engineering Design of Concentric Amphiphilic Microparticles for Spontaneous Formation of Picoliter to Nanoliter Droplet Volumes. Anal Chem 93, 2317–2326 (2021).

36. Wang, W., Zhang, M. J. & Chu, L. Y. Functional polymeric microparticles engineered from controllable microfluidic emulsions. Acc Chem Res 47, 373–384 (2014).

37. Li, W. et al. Microfluidic fabrication of microparticles for biomedical applications. Chemical Society Reviews vol. 47 5646–5683 Preprint at 10.1039/c7cs00263g (2018).

38. Rana, M., Ahmad, R. & Taylor, A. A microfluidic double emulsion plaoorm for spatiotemporal control of pH and particle synthesis. Lab Chip (2023) doi:10.1039/d3lc00711a.

39. Lee, S., De Rutte, J., DimaAeo, R., Koo, D. & Di Carlo, D. Scalable Fabrication and Use of 3D Structured Microparticles Spatially Functionalized with Biomolecules. ACS Nano 16, 38–49 (2022).

40. Keller, S., Hu, G. X., Gherghina-Tudor, M. I., Teora, S. P. & Wilson, D. A. A Microfluidic Tool for Fine-Tuning Motion of Sos Micromotors. Adv Funct Mater 29, (2019).

41. Liu, Q. et al. Self-Orienting Hydrogel Micro-Buckets as Novel Cell Carriers. Angewandte Chemie – International Edition 58, 547–551 (2019).

42. Yuan, H. et al. Phase-Separation-Induced Formation of Janus Droplets Based on Aqueous Two-Phase Systems. Macromol Chem Phys 218, (2017).

43. Ma, S. et al. Fabrication of microgel particles with complex shape via selective polymerization of aqueous two-phase systems. Small 8, 2356–2360 (2012).

44. Iqbal, M. et al. Aqueous two-phase system (ATPS): an overview and advances in its applications. Biological Procedures Online vol. 18 1–18 Preprint at 10.1186/s12575-016-0048-8 (2016).

45. Chao, Y. & Shum, H. C. Emerging aqueous two-phase systems: From fundamentals of interfaces to biomedical applications. Chemical Society Reviews vol. 49 114–142 Preprint at 10.1039/c9cs00466a (2020).

46. Kalra, A. P. et al. A Nanometric Probe of the Local Proton Concentration in Microtubule-Based Biophysical Systems. Nano LeJ 22, 517–523 (2022).

47. Thungon, P. D., et al. A Fluorescent Alcohol Biosensor Using a Simple microPAD Based Detection Scheme. Frontiers in Sensors 3, (2022).

48. Wang, H. et al. Ultrasensitive Picomolar Detection of Aqueous Acids in Microscale Fluorescent Droplets. ACS Sens 7, 245–252 (2022).

49. Dendukuri, D., Pregibon, D. C., Collins, J., Hatton, T. A. & Doyle, P. S. Continuous-flow lithography for high-throughput microparticle synthesis. Nat Mater 5, 365–369 (2006).

50. Teixeira, A. G. et al. Emerging Biotechnology Applications of Aqueous Two-Phase Systems. Adv Healthc Mater 7, (2018).

51. Ho, C. S., Kim, J. W. & Weitz, D. A. Microfluidic fabrication of monodisperse biocompatible and biodegradable polymersomes with controlled permeability. J Am Chem Soc 130, 9543–9549 (2008).

52. Krutkramelis, K., Xia, B. & Oakey, J. Monodisperse polyethylene glycol diacrylate hydrogel microsphere formation by oxygen-controlled photopolymerization in a microfluidic device. Lab Chip 16, 1457–1465 (2016).

53. Debroy, D., Oakey, J. & Li, D. Interfacially-mediated oxygen inhibition for precise and continuous poly(ethylene glycol) diacrylate (PEGDA) particle fabrication. J Colloid Interface Sci 510, 334–344 (2018).

54. Gruner, P. et al. Controlling molecular transport in minimal emulsions. Nat Commun 7, (2016).

